# Genetic diversity in facultatively sexual populations and its implications for the origins of self-incompatibility in algae and fungi

**DOI:** 10.1101/2021.04.04.438359

**Authors:** A.S.A. Smith, S. Penington, I. Letter, D.B. Wilson, G.W.A. Constable

## Abstract

The evolutionary mechanism that drove the establishment of self-incompatibility in early sexual eukaryotes is still a debated topic. While a number of competing hypotheses have been proposed, many have not received detailed theoretical attention. In particular, the hypothesis that self-incompatibility increases the benefits of genetic recombination in sexual haploids has been comparatively understudied. In this paper we address this topic by mathematically deriving how the probability of mating with a genetically distinct individual changes as a function of the presence or absence of self-incompatible mating type classes. We find that although populations with mating types successfully engage in sexual reproduction less frequently than their self-compatible competitors, they can nevertheless engage in useful sex with genetically distinct partners *more* frequently. This conclusion holds when the number of sexual reproductive events per generation is low (i.e. in small populations with low rates of facultative sexual reproduction). Finally we demonstrate the potential for frequency-dependent selection in competitive dynamics between self-compatible and self-incompatible types. These analytic results provide a baseline for studying the sex advantage enhancer model for the evolutionary origin of mating types within each specific hypothesis for the evolution of recombination.

**PACS:** 87.23.-n Ecology and evolution, 87.23.Kg Dynamics of evolution, 02.50.Ey Stochastic processes

**2000 MSC:** 37N25: Dynamical systems in biology, 60J70: Applications of diffusion theory (population genetics, absorption problems, etc.)

---

A primary challenge encountered by sexually reproducing organisms is that of locating a suitable mate (Lehtonen et al., 2012). We might therefore expect that any mutation that increases an organism’s ability to find a mate should be selected for, while any mutation that decreases an organism’s probability of encountering a mate should be selected against. Framed in this way, it is perhaps surprising that the vast majority of sexual organisms are not free to mate indiscriminately, but rather must find a complementary partner (Lehtonen et al., 2016).

Amongst most fungi, ciliates and algae, mating is restricted to occurring between individuals from distinct self-incompatible (SI) classes, termed mating types (Haag, 2007). These mating types are genetically determined by mating type loci that can be crudely understood as ancestral forms of the more familiar sex chromosomes present in many animals (Branco et al., 2017). Indeed, both empirical and theoretical studies support the view that mating types form an evolutionary precursor to the true sexes (Togashi et al., 2012; Lehtonen and Kokko, 2011). Unlike the true sexes however, which are defined by the morphological dissimilarity of their sperm-egg gametes (anisogamy), the gametes of distinct mating types are morphologically similar (isogamy).

The lack of size dimorphism between the sex cells of isogamous species raises a number of interesting evolutionary possibilities. For instance, in the absence of the disruptive selection that drives the evolution of the two sexes, the number of mating types is not restricted to two, with up to 23, 328 in fungi (Kothe, 1999) and 100 in ciliates (Phadke and Zufall, 2009). However, this morphological similarity also raises evolutionary questions. In anisogamous species, the evolutionary stability of self-incompatibility is intuitively clear; selection against sperm-sperm syngamy is driven by low viability of the resultant ill-provisioned zygote while selection for egg-egg syngamy is weakened by their low encounter rate. With such factors not present in isogamous species, we can then ask if self-incompatibility can remain evolutionarily stable in these systems and, more fundamentally, how it even first arose.

Most models of the early evolution of sexual reproduction suppose the existence of a “unisexual” early ancestor that mated indiscriminately and from which mating type self-incompatibility systems evolved (Heitman, 2015). However, there is as yet no empirical evidence indicating that such a unisexual system preceded the mating type self-incompatibility systems in isogamous species. Therefore if this supposition is true (and this seems likely given the difficulty of evolving mechanisms for sexual reproduction and two self-incompatibility systems simultaneously) the rapid loss of this unisexual ancestor indicates a nascent advantage to self-incompatibility in the evolution of early sex. We next turn our attention to the stability of isogamous self-incompatibility systems.

The loss of self-incompatibility has occurred in multiple isogamous species, with phylogenetic analyses showing that self-compatibility is a derived trait that has evolved on top of preexisting mating type self-incompatibility systems (Roach et al., 2014). This is termed homothalism. Homothalism comes in many distinct forms (Wilson et al., 2015) including: primary homothalism (individuals posses multiple ancestral mating type alleles on the same chromosome), same-sex mixing (individuals of the same mating type can mate) and mating type switching (individuals carry multiple mating type alleles with alternate mating types expressed in alternate generations of asexual progeny). However, the effect of each of these mechanisms is to remove the barriers preventing the progeny of a parent (or recent ancestor in the case of mating type switching) from mating with each other. It is clear that the evolutionary advantage of homothalism is the improved chance of finding a mate for sexual reproduction. This can provide reproductive assurance for obligately sexual species or facultatively sexual species in which sexual reproduction is associated with the formation of a resistant state. However, in order to understand when self-incompatibility is selected against we must not just identify the costs of self-incompatibility, but also its benefits.

Multiple theories have been proposed as to the evolutionary pressures that may have driven the evolution of self-incompatibility in early sexual eukaryotic ancestors (reviewed in (Billiard et al., 2011)). Some of these assume a physiological advantage to SI mating types. These include the “by-product model” (that a bipolar pheromone-receptor recognition system improves the efficiency of syngamy between gametes (Hoekstra, 1982; Hadjivasiliou et al., 2015; Hadjivasiliou and Pomiankowski, 2019)) and the “ploidy model” (that self-incompatibility brings together distinct mating type alleles that in concert may signal a diploid form, with which syngamy should not be instigated (Haag, 2007)). Other models suggest that genetic conflict may lie at the heart of the evolution of self-incompatibility. These include the “organelle inheritance model” (that SI alleles in fact encode for donor-receiver roles in the uniparental inheritance of cytoplasm (Hurst and Hamilton, 1992; Hadjivasiliou et al., 2012)) and the “selfish genetic element model” (that SI alleles emerged as selfish genetic elements that evolved to target “uninfected” non-self gametes for mating). A final class of models, and the focus of interest for this paper, investigates how the costs and benefits to sex and genetic recombination may be respectively suppressed or amplified by self-incompatibility.

The “inbreeding depression avoidance model” suggests that as self-incompatibility promotes disassortative mating, it reduces the rate at which rare deleterious mutations become homozygous in inbred offspring and may therefore be selected for (Charlesworth and Charlesworth, 1979; Uyenoyama, 1988a,b). However, as this hypothesis relies on species being diploid, and with sporophytic self-incompatibility determined in this diploid stage, it is inappropriate for isogamous species which are mostly haploid with gametophytic selfincompatibility determined at the haploid stage (Billiard et al., 2011). (While the model is appropriate for SI alleles in plants, in this scenario self-incompatibility has evolved on top of a pre-existing anisogamous mating system and therefore lies outside the scope of this paper.) In contrast, the “sex advantage enhancer model”, which suggests that self-incompatibility amplifies the benefits of genetic recombination, is suitable for haploid isogamous species. However, with rare exceptions (Nauta and Hoekstra, 1992; Czaran and Hoekstra, 2004), it has received little theoretical attention.

From a modelling perspective, the reason that the sex advantage enhancer model has been tackled infrequently is clear. The hypothesis is straightforward to formulate verbally: while the benefits of recombination in haploids cannot be realised if syngamy occurs between identical haploid clones, the probability that this occurs should be decreased by the presence of SI alleles in the population. However, it is significantly harder to formulate the hypothesis mathematically; the precise mechanisms that drive selection for genetic recombination are themselves still an intensely debated subject (Agrawal, 2006). A complete investigation of the feasibility of this hypothesis therefore requires that the effects of selfincompatibility are tested within each recombination hypothesis framework (e.g. Fisher–Muller hypothesis, the Hill–Robertson effect, the Red Queen hypothesis (Hartfield and Keightley, 2012)), which themselves only yield selection for recombination under a limited range of scenarios.

An alternative approach is to abandon any attempt to model selection for recombination and instead take a coarser-grained perspective. For instance, one could impose an advantage to non-self mating ‘by hand’, such that the progeny of SI parents have an enhanced reproductive rate (Nauta and Hoekstra, 1992; Czaran and Hoekstra, 2004). However, this leads to a rather circular argument (self-incompatibility is selected for if the reproductive rate enhancement exceeds the cost of mate finding) and does not capture the feedback between sexual reproduction, self-incompatibility and population diversity.

In this paper we take a simplified view that neglects specific hypotheses for the benefits of recombination, but nevertheless cuts to the core of the verbal formulation of the sex advantage enhancer model. In particular, by investigating the relationship between population-level genetic diversity and SI systems, we calculate how the probability of mating with a genetically distinct individual varies as a function of the presence or absence of mating type alleles and depending on their number. We also investigate how these factors interplay with the rate of facultative sex in the population, which recent theoretical studies have suggested may play a role in the evolution of mating type number (Constable and Kokko, 2018; Czuppon and Constable, 2019; Krumbeck et al., 2020; Berríos-Caro et al., 2021). In the context of models of the evolutionary origins of self-incompatibility, considering the role of facultative sex is important as it is highly likely that sexual reproduction was rare in its early evolutionary history.

## 1. The model

We consider a haploid population in which the mating type of an individual is determined by one of an infinite set of potential alleles at a single locus. We shall assume that the genome is long and this locus is small so that we may approximate its position by a single point *m* within the normalized genome length [0, 1] (see Figure 1). Using a Moran modelling framework in which birth and death events are coupled, the population is fixed to a constant size *N*. Each individual will enter a reproductive phase at rate *N*; this time scaling will be convenient for our calculations.

**Figure 1:**
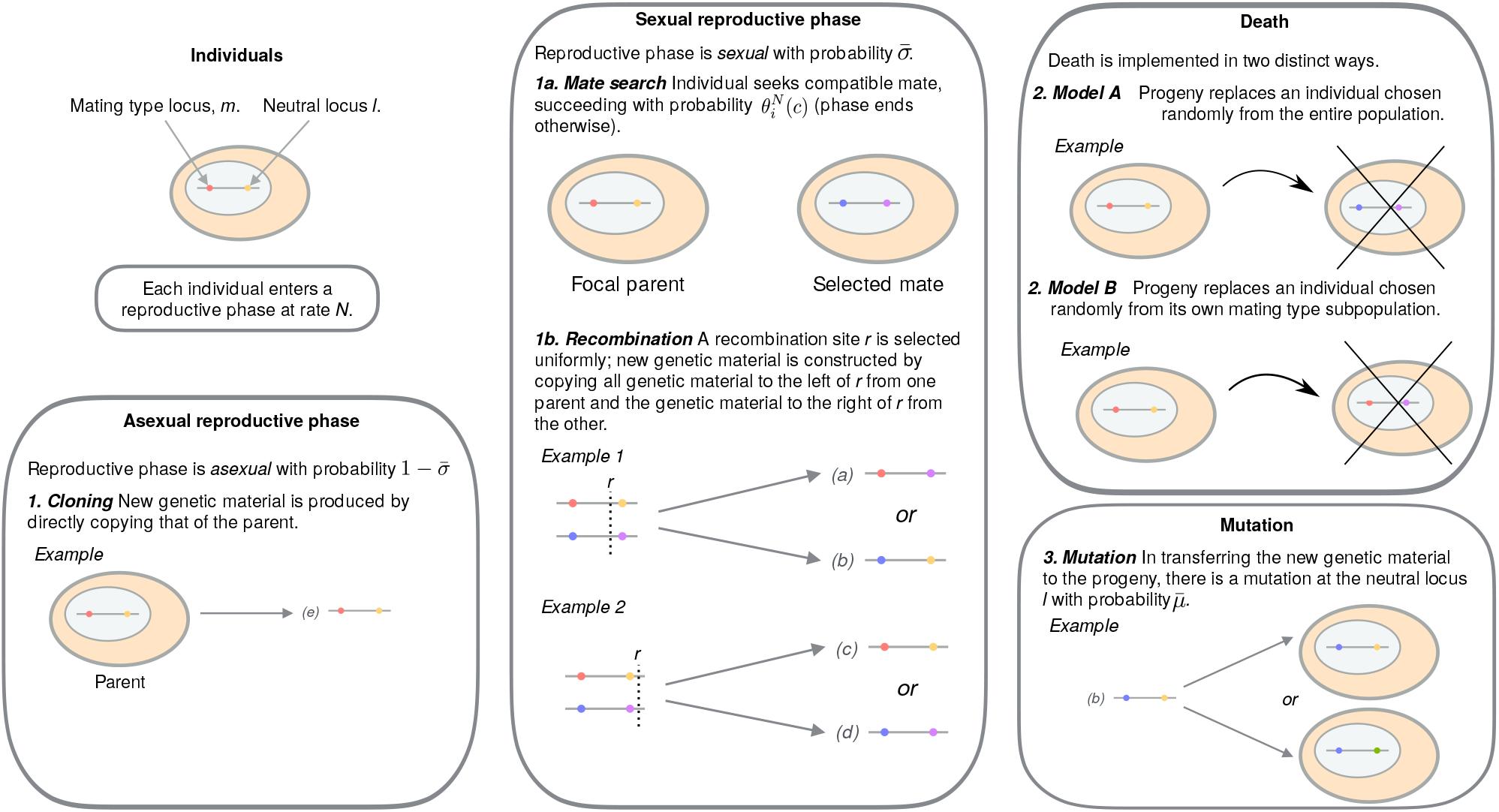
Figure illustrating model dynamics.

For some fixed parameter 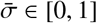, we suppose that each individual reproduces asexually at rate 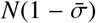. Upon being chosen for an asexual reproduction event, an individual is created that inherits its parental mating type. Meanwhile, for each individual, at a rate 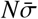 another individual is randomly chosen to attempt to instigate a sexual reproduction event. Here 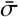 approximately captures the ratio of rates of sexual to asexual reproduction. For facultatively sexual populations 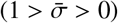, we will consider three scenarios for the mating dynamics.

Firstly, we will consider a ‘unisexual’ population in which the chosen individual can mate with any other individual in the population irrespective of mating type. The probability that this individual can successfully mate with another chosen randomly from the population is thus one. (We choose the terminology unisexual to emphasise that here we may be capturing a sexual form that preceded mating type self-incompatibility (Roach et al., 2014).)

Secondly, we consider a population with *k* self-incompatible (SI) mating types. We denote by *N*_*i*_ the number of individuals of mating type *i*, such that 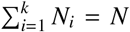. Having been chosen to attempt a sexual reproduction event, an individual begins what we term a *mating round* (described below) during which they are given a number of attempts to randomly select a partner carrying a distinct mating type allele from the population.

Finally we consider a case with *k*−1 SI mating types and a single unisexual (self-compatible) type. If an individual from one of the first *k*−1 SI types is chosen for sexual reproduction, it instigates a mating round during which it attempts to find a suitable mate. Meanwhile if an individual from the unisexual type is chosen for sexual reproduction, it finds a mate with probability one.

### 1.1. Mating rounds

SI mating types are by definition unable to mate with those of the same mating type. They are however permitted a random number of attempts to find a compatible mate. We call the individual which attempts to find a mate the *focal parent* (see Figure 1). During each attempt, a potential mate is chosen uniformly at random from the *N*−1 remaining individuals in the population. The attempt is rejected if the chosen mate is of the same mating type and accepted otherwise. The mating round ends either once all allowed attempts are rejected or after the first reproduction with a compatible partner. (Individuals thus sexually reproduce at most once per mating round.)

The number of attempts an individual makes will be random, and we will assume a Gibbs-type distribution for the number of attempts: for a fixed parameter *c >* 0, if *A* is the random number of attempts made, we will take *P*[*A* = *n*] ∝*e*^−*cn*^ for all *n* ≥1 and *P*[*A* = *n*] = 0 otherwise. In particular, this implies that *A* is geometrically distributed with parameter 1 − *e*^−*c*^ and

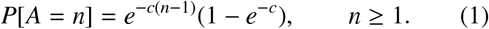

We note that the limit *c*→∞ corresponds to each individual having exactly one chance to find a compatible mate (i.e. a large cost to mate finding) and the limit *c*→0 corresponds to the case where individuals may make as many attempts as necessary to find a compatible mate (i.e. no cost to mate finding). The parameter *c* can thus be understood as interpolating between mass-action encounter rates at one extreme (*c* → ∞) and active, highly effective mate searching on the other (*c* → 0).

For fixed *c >* 0, an individual of SI mating type *i* mates with an individual of mating type *j* ≠ *i* with probability

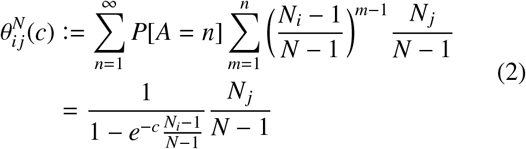

and thus successfully finds a compatible mate during a mating round with probability

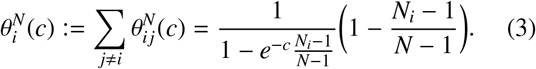

### 1.2. Recombination

If the individual is successful in finding a compatible mate during the mating round, the juvenile inherits genetic material from both parents as follows. A recombination site *r* is selected uniformly at random on the unit interval [0, 1] and the offspring inherits all genetic material to the left of *r* from one parent and the genetic material to the right of *r* from the other (see Figure 1).

We consider a highly polymorphic neutral genetic marker at position *l* ∈ [0, 1] on the genome. It is one of an infinite set of potential alleles denoted by elements in the set {*a*_1_, *a*_2_, …} The allelic type of an individual at *l* is assumed to have no influence on its mating type. We will denote by *d* = |*m*−*l*| the distance on the genome between the locus determining the individual’s mating type and the locus determining its allelic class.

In a sexual reproduction event, both mating and allelic types may be inherited from either parent or they may individually be inherited from distinct parents. As the recombination site is chosen uniformly on the interval [0, 1] and the loci carrying the mating type and allelic type are at distance *d* apart, the probability that these two traits are inherited from distinct parents is *d*. The probability of each possible outcome can be tabulated as in Table 1.

**Table 1:**
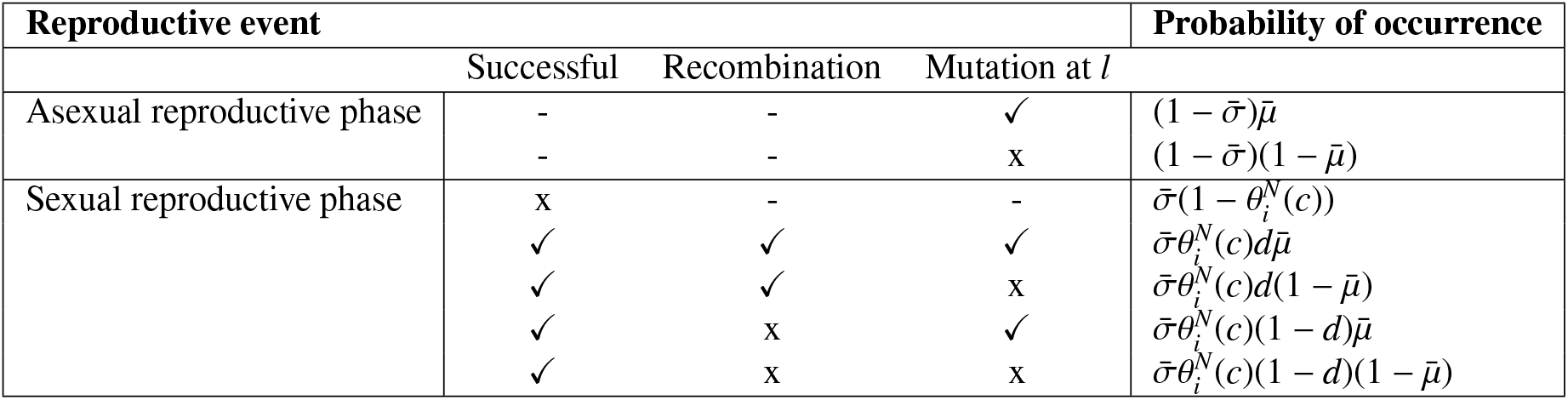
Inheritance table showing possible outcomes and their probabilities when an individual of the *i*th mating type enters a reproductive phase.

We note that the assumption that there is only one recombination site significantly simplifies our analysis. In the case where we consider genetic diversity on a separate chromosome, we set *d* = 1*/*2 (i.e. there is no linkage between the mating type allele and the neutral marker).

### 1.3. Mutation

In addition we will suppose that in the case of either sexual or asexual reproduction, the genetic material at the neutral locus *l* is incorrectly transmitted to the progeny with probability 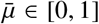. It is assumed that the mutation is to a selectively neutral allelic type not yet seen in the population.

### 1.4. Replacement

In asexual and sexual reproduction events, the new individual replaces another individual in the population. We will consider two different scenarios: in Model A, the individual that is replaced is chosen uniformly at random from the whole population, and in Model B, the individual is chosen uniformly at random from the subpopulation of the mating type of the new individual. (Note that in Model B, the size of the mating type subpopulations is constant.)

## 2. Mathematical analysis

### 2.1. Deterministic dynamics for SI allele frequencies

We begin by briefly discussing the costs to mating types in terms of mating opportunities and how they manifest within the context of our model. As such we will focus here only on the dynamics of the mating type alleles under Model A (that is, we allow the frequencies of mating type alleles to vary through time).

Firstly we note that when the population is entirely asexual 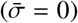, the model reduces to a neutral Moran model. In this case the mating type alleles do not experience any selective pressure, and eventually all but one of the alleles will be lost due to genetic drift. If a unisexual type is introduced into this asexual population it will drift to either fixation or extinction (the per-capita rate of reproduction is 1 for both asexual and unisexuals in this scenario). We refrain from attaching any biological importance to this neutrality, as in terms of reproductive opportunities benefits to sexuality (where sex is associated with the formation of an environmentally resistant form) or costs (sexual reproduction often incurs a larger time cost) could be accounted for in a more general model. As we are primarily interested in the transition from unisexuality to mating types in the current study, we do not account for these behaviours here.

For a population with *k* sexually reproducing self-incompatible mating types 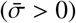, the dynamics of the mating type frequencies *p*_*i*_ = *N*_*i*_*/N* are given by

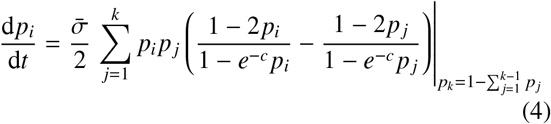

in the infinite population size (deterministic) limit (see Appendix A). In the limit of mass-action encounter rate dynamics (*c* → ∞) the equations revert to those explored in (Constable and Kokko, 2018; Czuppon and Constable, 2019).

For general *c >* 0, the population approaches an equilibrium in which all mating types are equally represented (the system has a single attracting fixed point at *p*_*i*_ = 1*/k* for all *i*). However, the stability of this fixed point, measured by the eigenvalues *λ*_*i*_, decreases with decreasing *c* and increasing *k* (which both lead to an increased probability of finding a mate) and decreasing 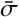;

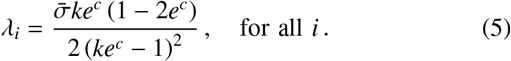

Following a mutation event that introduces a new type (*k* → *k* + 1), the fixed point (now with *p*_*k*+1_ ≈ 0) becomes unstable, while the fixed point at *p*_*i*_ = 1*/*(*k* + 1) is stable, with eigenvalues given by Eq. (5) (with *k* terms replaced by *k* + 1). This means that there is always selection for a larger number of mating types. However, the strength of selection for these new mating types decreases as *c* and 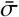 decrease, and as *k* increases. The decrease in selection strength in these parameter regimes will lead to higher rates of extinction of mating type alleles when genetic drift is accounted for (Czuppon and Rogers, 2019). This has been previously quantified analytically in the limit of *c* → ∞ (Constable and Kokko, 2018; Czuppon and Constable, 2019).

We have seen that as *c* approaches zero (i.e. each individual has an infinite number of attempts to mate) the reproductive advantage to rare types is reduced. However, perhaps surprisingly, this advantage is never eliminated entirely. This is because although a rare type has the same per capita probability of finding a partner as more common types, it in fact has a higher per capita probability of being found (see Appendix A).

For a population with (*k* − 1) sexually reproducing SI mating types 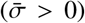, and a single unisexual type (frequency 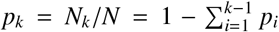) the equivalent dynamics (see Appendix A) are given by

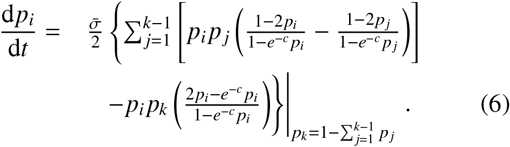

For general 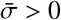, the unisexual type is selected for (the system has an attracting equilibrium at *p*_*k*_ = 1). When the unisexual type is resident and SI mating types are rare (*p*_*k*_ ≈1), selection against the SI types is weak due to the low probability of encountering another SI type. (In fact the eigenvalues of the fixed point *p*_*k*_ = 1 are zero, indicating neutral stability at leading order.) Nevertheless when SI mating types reach an appreciable frequency they will be selected against, ensuring the success of unisexuals over long timescales.

In this section (using Model A) we have seen that the frequencies of each mating type class lie in the region 1*/k* for a population of *k* SI classes. In the following sections we will therefore assume for tractability that each mating type has precisely this frequency by coupling birth and death events within a mating-type class (see Model B). However, we have also seen in this section (using Model A) that all *k* SI mating types are driven to extinction upon the introduction of a unisexual type. To tackle this scenario in later sections, we will also often couple birth and death events within the unisexual class (in addition to the mating type classes). While this coupling is clearly less well-justified for the case of unisexual invasion, we will see that this can be understood as assuming that the unisexual invasion occurs slowly.

### 2.2. Stochastic dynamics of neutral markers

Having looked at the dynamics of the mating type frequencies, here we focus on the dynamics of the neutral markers under the different mating systems. From now on we consider Model B, i.e. we assume that the sizes of the mating type subpopulations are fixed. We aim to calculate the probability that two individuals share the same neutral genetic marker when randomly sampled from an asexual population, a unisexual population and a population with *k* SI mating types respectively. We then proceed to approximate this same quantity for a mixed population with a single unisexual type and *k*−1 SI types. In the following subsections we develop the mathematical machinery required for this calculation. Readers less interested in the specific mathematical details may proceed to the Results section.

#### 2.2.1. The rare sex, rare mutation regime

Our initial interest lies in the case that both sex, and mutations at the locus *l* are rare; in particular, it will be assumed that for some parameters *σ >* 0 and *µ >* 0,

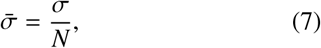

and

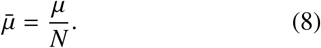

The parameter *σ* may be interpreted as the average number of initiated sexual mating rounds per *N* reproductive events. The parameter *µ* may be interpreted as the average number of successful reproductive events which terminate with a mutation per *N* successful reproductive events. We stress that this is per *successful* reproductive event, noting that an individual that instigates a sexual reproductive event may not be successful in the production of an offspring if a compatible mate is not found during the mating round. Moreover, as the average number of reproductive events per unit time varies with the relative rate of sexual reproduction (from *N* in the asexual 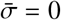 case to 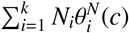 in the obligately sexual 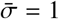 case) so too does the number of mutations per *N* reproductive events. This leads to a natural definition of the effective rate of mutation given by

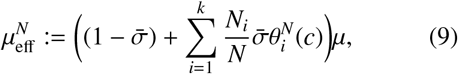

which is the rate per individual of introducing a new mutation to the population. Equivalently, 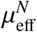 is the average number of mutations per *N* reproductive phases.

In the same vein, we may define an effective recombination rate

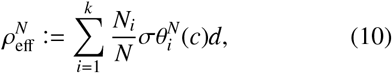

that is, the average per individual rate of a crossover between the mating type allele and the allele at the neutral marker. Equivalently, 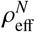 is the average number of crossovers between these same alleles per *N* reproductive events. The quantity 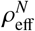 measures the degree of gene flow between SI mating type subpopulations.

#### 2.2.2. Diffusion approximation of allele frequencies at the neutral marker

In the rare sex, rare mutation regime (as described in the previous section), as *N* → ∞ the allele frequencies within mating type are well approximated by a *k*-dimensional diffusion process which can be seen as a solution to a stochastic differential equation (SDE). A brief account of diffusion processes and how they may be used to approximate continuous-time Markov chains is given in Appendix B.

Recall that the allelic type at the neutral locus *l* is one of an infinite set of potential alleles denoted by elements in the set {*a*_1_, *a*_2_,*…*}. For the reference type, *a*_1_, denote for each *i* = 1, *…, k*,

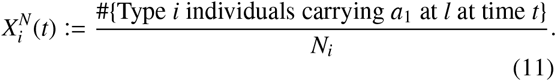

That is, 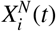 denotes the proportion of individuals of the *i*th mating type which carry the allele *a*_1_ at the neutral locus *l* at time *t*. Passing to the large population limit (*N* → ∞), we further define *p*_*i*_ as the fraction of individuals of mating type *i* (i.e. *N*_*i*_*/N* →*p*_*i*_) and *x*_*i*_ as the initial frequency of *a*_1_ in the *i*th mating type class (i.e.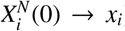). In Appendix F we then show that *X*^*N*^ converges to the solution of the system of stochastic differential equations given by

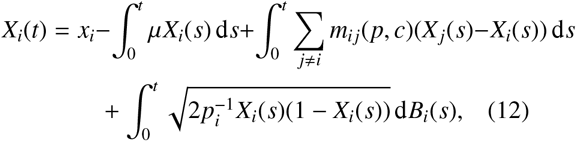

where, for each *i, j* =1, *…, k, B*_*i*_ is an independent Brownian motion and

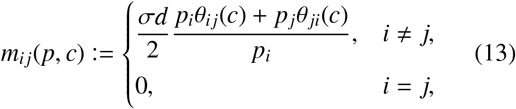

where 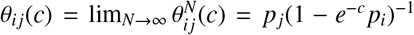. Note that we can write

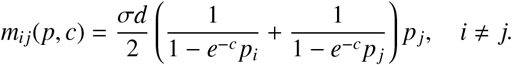

#### 2.2.3. Approximate genealogies

By recording only the proportion of each subpopulation carrying the reference type *a*_1_ at the neutral marker, *X*_*i*_(*t*), we do not retain information about how individuals are related to one another. However, looking backwards-in-time, it is possible to describe the genealogical trees relating individuals in a random sample from the population.

We wish to establish the genealogy at the neutral marker *l* of a finite sample of individuals taken from the population *X*^*N*^. Denote by 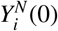 the number from this sample of mating type *I* We trace the ancestry of the neutral marker at the neutral locus, denoting by 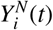 the number of individuals of the *i*th mating type that are ancestors to the original sample *t* units of time into the past.

Passing to the large population limit *N* → ∞,it is shown in Appendix G that the process 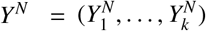 approximates a structured coalescent process *Y* which evolves as follows. If *Y*(*t*) = (*y*_1_,*…,y*_*k*_) then

1. each pair of type *i* ancestral lines *coalesces* at rate 2*/p*_*i*_, independently of one another; i.e. *y*_*i*_ ↦ *y*_*i*_ −1 at rate

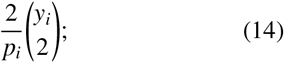
2. each ancestral line of type *i* becomes an ancestral line of type *j* at rate *m*_*i j*_(*p, c*), independently of one another; i.e. (*y*_*i*_, *y* _*j*_)↦ (*y*_*i*_ − 1, *y* _*j*_ + 1) at rate

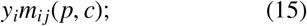

and
3. each ancestral line experiences a mutation at total rate *µ*, independently of one another. (There is no change to the number (or mating types) of individuals that are ancestors to the original sample during this event.)

This result is proved in Appendix G.

The parameter *m*_*i j*_(*p, c*) may thus be thought of as an effective per individual rate of a crossover between a type *i* and type *j* individual. Models for genealogies of this type are often referred to as structured coalescents, or island models (*c.f*. Durrett (2008)) and, since the recombination events (item 2 above) manifest as an effective migration of ancestors between mating type subpopulations, we will usually refer to such events as a “migration”, for consistency with the island model. An illustration of this process is depicted in Figure 2.

**Figure 2:**
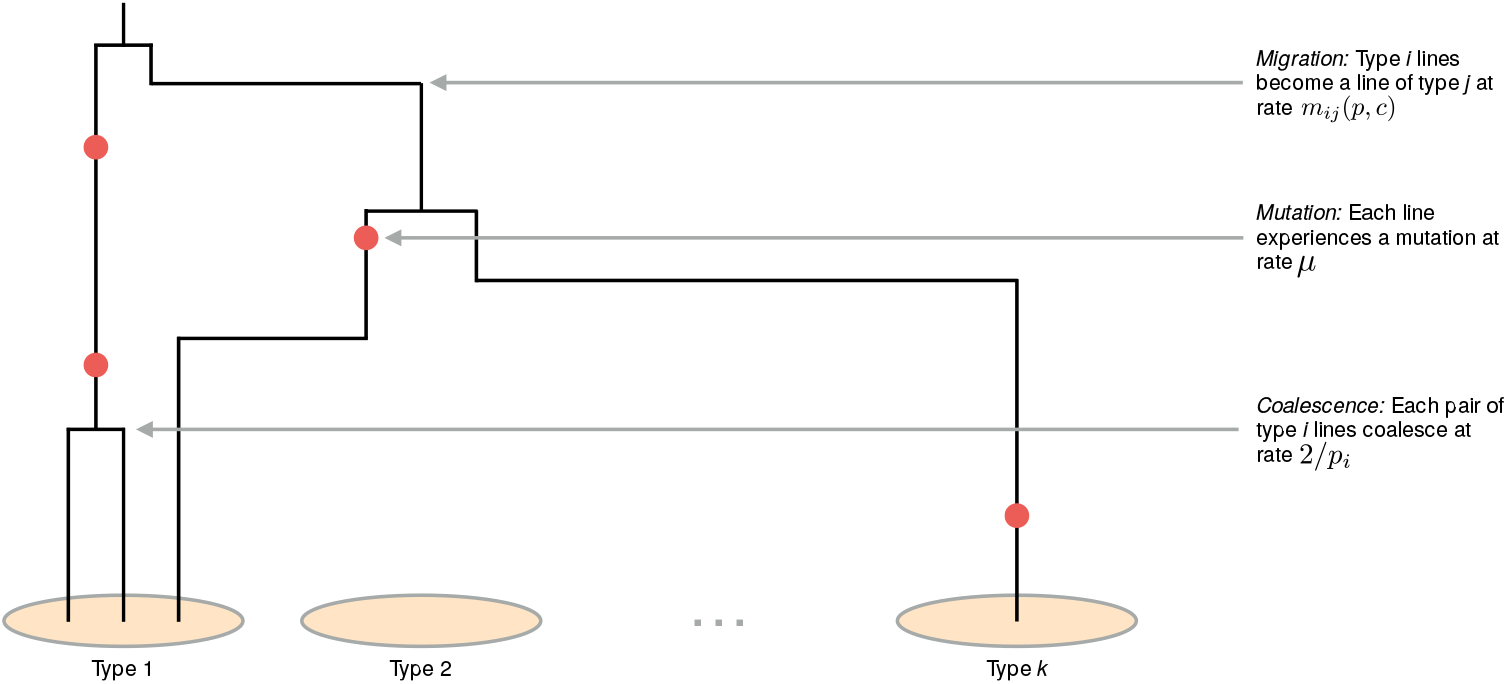
Figure illustrating the dynamics of the genealogical process *Y*(*t*).

In Appendix E, we demonstrate that the processes *X* and *Y* are dual in the sense that given *X*(0) = *x* and *Y*(0) = *y*,

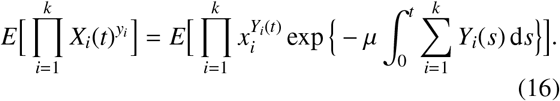

Relations of this kind are frequently of interest in the study of stochastic processes (see Appendix D). In Appendix E, we make explicit use of this duality in proving the uniqueness of solutions to the system of Eqs. (12).

#### 2.2.4. Probability of identity by descent: monomorphic populations

It is clear from our description of the genealogical process that two individuals sampled from the population will have a common ancestor if we look far enough back into the past. We will say that the two individuals are *identical by descent* if they are of the same allelic type at the locus *l*; this will be the case if neither of the sampled individuals experience a mutation before their eventual coalescence. Let us define *π*_*i j*_ to be the probability that an individual sampled from the type *i* mating type subpopulation and an individual sampled from the type *j* subpopulation (distinct from the first if *i* = *j*) are identical by descent, i.e. the probability that they carry the same allelic type at the neutral marker *l*.

We begin by considering a population of *k* self-incompatible mating types. (We will see in the Results section that cases of monomorphic asexual populations and unisexual populations can be obtained as limiting forms of this scenario.) In Appendix H, we derive recursions for the probabilities *π*_*i j*_, *i, j* = 1, 2, *…, k*; how-ever, we are not able to find tractable solutions in this generality. In Section 2.1, we saw that demographic noise leads to fluctuations in *N*_*k*_ which can cause extinctions, but as we are interested in the behaviour of the population before such extinctions occur, we begin by fixing *p*_*i*_ = 1*/k* for all *i*. This approach has previously been taken by (Ennos and Hu, 2019), and in the case of *k* SI types is equivalent to working in the regime *k/N* ≪1 and *c >* 0, which is a biologically reasonable regime for scenarios in which mating type extinctions are rare (Czuppon and Rogers, 2019).

The above simplification induces some additional symmetry to the system and increases the analytic tractability of the problem. In this setting, only two quantities are of interest; they are *π*_*d*_, the probability that two individuals sampled from distinct mating type subpopulations are identical by descent, and *π*_*s*_, the probability that two individuals sampled from the same subpopulation are identical by descent. Observe that *π*_*ij*_ ≡ *π*_12_ =: *π*_*d*_ if *i* ≠ *j* and *π*_*ij*_ ≡ *π*_11_ =: *π*_*s*_ if *i* = *j*. Similarly, we can obtain an expression for 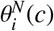 (see Eq. (3)) in this limit:

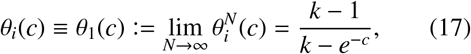

and for *i* ≠ *j*,

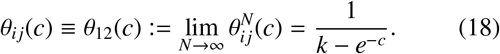

We can now define the “migration” rate, *m*, (see Eq. (13)), which is now equal across types, by

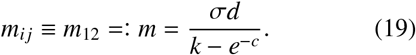

Recall that by “migration” we are referring to the effective migration of lineages between subpopulations corresponding to gene flow between types resulting from recombination.

In Appendix H it is shown that

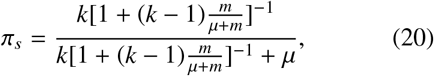

and

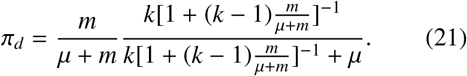

We similarly define *π*_tot_ to be the probability that two distinct individuals, chosen uniformly at random from the whole population, are identical by descent, i.e. carry the same allele at the neutral marker *l*. Since the pair are of the same mating type with probability 1*/k* and of distinct types with probability (*k* − 1)*/k*, the quantity *π*_tot_ is given by the weighted average of *π*_*s*_ and *π*_*d*_,

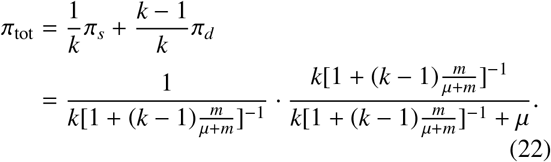

We next consider a unisexual population. For a unisexual population, there is only a single well-mixed population (*k* = 1). Importantly, while sexual reproduction is now possible between all individuals, this in fact has no impact on the probability of identity by descent at the neutral locus. Without the population structure imposed by self-incompatibility, the inheritance of neutral alleles during sexual reproduction is effectively the same as that under asexual reproduction. We thus find that the probability of identity by descent in the unisexual population is given by

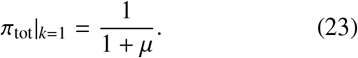

#### 2.2.5. Probability of identity by descent: mixed populations

We next consider a mixed population consisting of *k* − 1 self-incompatible mating types and a single unisexual type. We have seen previously that in the absence of any costs to unisexuality, the unisexual type always sweeps to fixation. We therefore consider three scenarios under which *π*_*i j*_ might be calculated. In the first scenario the frequency of the unisexual type is held at a constant frequency with respect to the *k* − 1 SI types (each at equal frequency). In the second, a rare unisexual type arises in a population of *k* − 1 SI types at equilibrium. In the third, *k* − 1 rare SI types arise in a population of the unisexual type at equilibrium.

We first consider the setting of a population subdivided into *k* allelic classes, one of which is unisexual. We will denote by *u* ∈ {1, 2, …, *k*}the index of the unisexual sub-population. For the sake of clarity, in this revised model individuals of type *u* are able to mate indiscriminately, and so always find a compatible mate, and individuals of type *i* ≠ *u* may mate only with those belonging to a class distinct from that of themselves.

In this setting, the probabilities 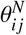, as computed in Eq. (2), are now given by

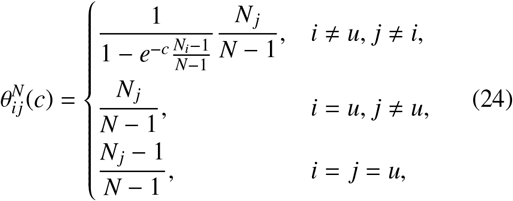

converging, as *N* → ∞, to

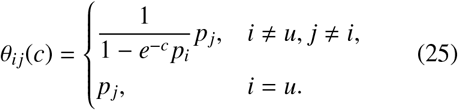

(Here, as above, *p*_*i*_ = lim_*N*→∞_ *N* _*i*_*/N*.) Following the mathematical reasoning and methodology above, including the proofs in Appendix E and Appendix F, it can be established that the proportion of type *i* individuals carrying the reference type *a*_1_ at the neutral marker *l* converges to the system of stochastic differential equations defined in Eq. (12), with the “migration” rates *m*_*i j*_ as defined in Eq. (13) but with the values of *θ*_*i j*_ given by the new values in Eq. (25). To be explicit, the migration rates become

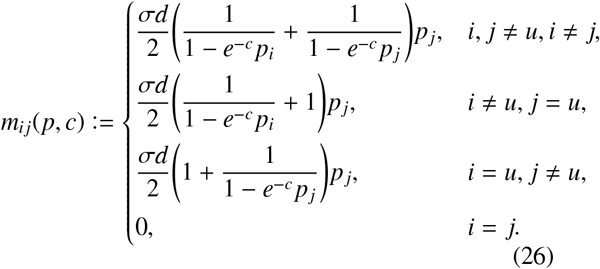

Similarly, the backwards-in-time genealogical process converges to the same structured coalescent as above but with these migration rates.

From this limiting genealogical process we wish to compute the probabilities of identity by descent *π*_*i j*_, but again find that while theoretically possible, it is mathematically impractical to do so without imposing some further symmetry. We will assume that a proportion *α* ∈ (0, 1) belong to the unisexual class, *u*, and so set *p*_*u*_ = *α*. We additionally assume that the remaining classes are held in constant frequency, setting *p*_*i*_ = (1− *α*)*/*(*k* − 1) for all *i* ≠ *u*. In this case we may write

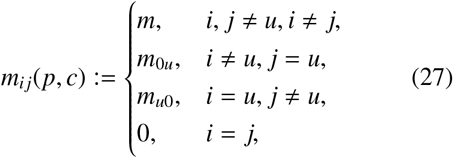

where

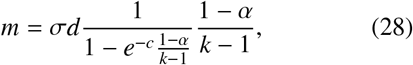

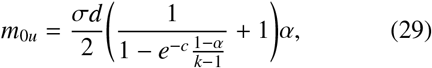

and

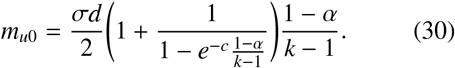

These migration rates are depicted in Figure 3.

**Figure 3:**
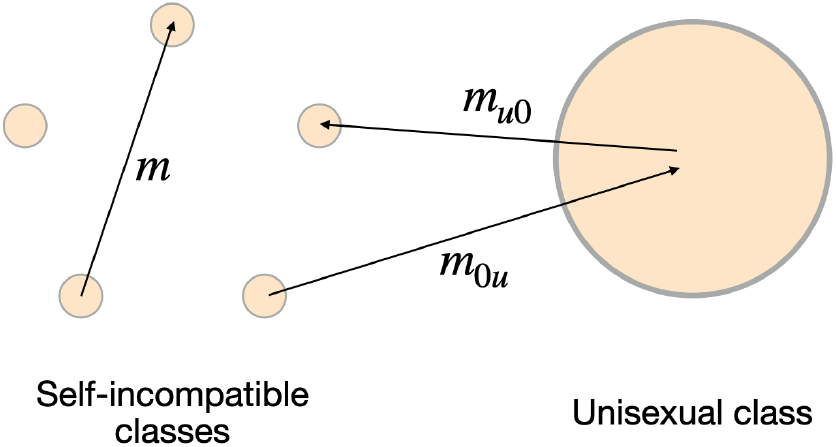
Migration rates between mating type classes in the structured coalescent for a population composed of both self-incompatible and unisexual types

Let us denote by *π*_*s*_ the probability that two individuals chosen from the same SI class are identical by descent; by *π*_*d*_ the corresponding probability for two individuals chosen from distinct SI classes; by 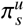 again the corresponding probability for two individuals both of the unisexual class; and by 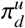 the corresponding probability for two individuals, one of the unisexual class and one of an SI class. A system of Eqs. (I.1-I.4) for the probabilities 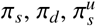 and 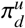 is derived in Appendix I.

Though straightforward to evaluate numerically, the resulting equations are too algebraically complex to be worthwhile reproducing here. Tractable analytic solutions are available in the case *α* = 1*/k*, a case of interest in our simulations, and further, in the (somewhat trvial) case that *c* = ∞. In this setting *m* = *m*_0*u*_ = *m*_*u*0_ = *σd/k* and two lineages within the same class coalesce at rate 2*k*, independent of whether this class is self-incompatible or unisexual. Because of this symmetry, 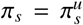 and 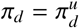 and the system of Eqs. (I.1-I.4) reduces to that of a population consisting of *k* self-incompatible types held in constant frequency (with *c* → ∞). In particular, in this special case,

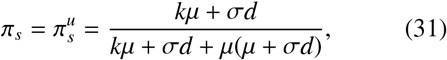

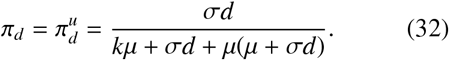

Suppose next that we have a single unisexual mutant emerge in an SI population with *k* − 1 mating types, leading to *k* types (including the unisexual type) in the population in total. We modify our Model B in this case to allow the size of the unisexual subpopulation to change; when a new individual is of the unisexual mating type, it replaces an individual chosen uniformly at random from the whole population. Initially the population is in the equilibrium described by Eqns. (20), (21), (22) with *k* replaced by *k* − 1, but otherwise unaffected by the presence of the rare mutant. Initially the mutant subpopulation will be fixed for a single neutral marker, and so the probability of identity by descent (IBD) when mating within its own subpopulation is one. Meanwhile when picking a partner from its ancestral mating type subpopulation (with probability 1*/*(*k* − 1)) it inherits the same probability of IBD as within that ancestral population. This leads to the following expressions for the probability of IBD for individuals sampled from the same (*π*_*s*_ and 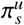) or distinct (*π*_*d*_ and 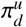) subpopulations, for SI and and unisexual types respectively:

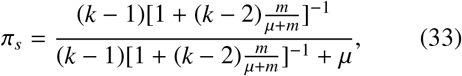

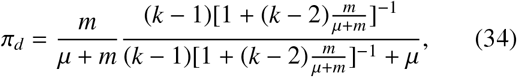

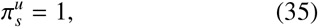

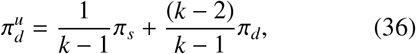

where *m* = *σd*(*k* − 1 − *e*^−*c*^)^−1^ is taken from Eq. (19).

Finally, we envisage a scenario in which (*k* − 1) mutant SI types emerge in a unisexual population at equilibrium. We note that this is a rather artificial scenario (except in the case *k* = 2), in the sense that it is unlikely that multiple SI mating types will evolve simultaneously. Nevertheless we consider this case as the natural symmetric situation to that discussed previously. The probability of IBD for individuals sampled from within the unisexual subpopulation will be unchanged by the presence of these rare mutants (i.e. will still be given by Eq. (23)), a property that will also be inherited by individuals sampled between the ancestral unisexual population and mutant SI populations. In the unlikely event that two individuals are sampled from within a given SI subpopulation, the probability of IBD is one. This leads to the following expressions for the probability of IBD:

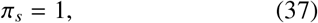

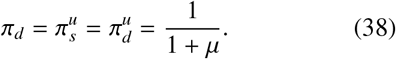

## 3. Results

### 3.1. Genetic diversity

Nei (1973) defines the *heterozygosity* or *gene diversity* of a population at a fixed locus *l* to be the probability that two individuals sampled uniformly at random carry distinct alleles at *l*. Since we are considering a haploid population, we prefer the term gene diversity and will use this exclusively henceforth. Nei’s measure of diversity, which we denote by *h*_tot_ and *h*_sub_ for the total population and mating type subpopulations respectively, can be expressed as *h*_sub_ = 1 − *π*_*s*_ and *h*_tot_ = 1 − *π*_tot_. We proceed to consider these calculations in scenarios of particular interest, focusing on how genetic diversity within the monomorphic populations changes as a function of the population parameters.

#### 3.1.1. Purely asexual population

The first case of interest, which will be used as a basis for comparison, is that of a purely asexually reproducing population. This situation is recovered by taking *σ* = 0 in Eqs. (19) and (22).

The case *k* = 1 corresponds to the whole population reproducing according to Moran model dynamics and the subsequent genealogical process being given by the unstructured Kingman coalescent. In this case, *π*_*d*_ has no meaning since the single ‘subpopulation’ and the metapopulation coincide. The probability of identity is given by

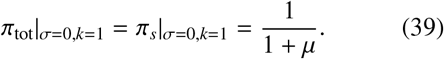

Thus Nei’s gene diversity, *h*_tot_ = 1 − *π*_tot_ increases as a function of mutation probability, 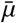, and population size, *N*, where we recall from Eq. (8) that 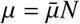.

While mating type classes are lost when *σ* = 0 and the sizes of the subpopulations are allowed to change, it is nonetheless useful as a basis of comparison to consider the above quantities when the mating type frequencies are held constant artificially at 1*/k*. In this case, the population is reduced to a set of *k* non-interacting asexually reproducing subpopulations, and

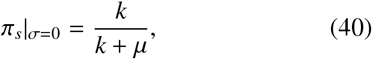

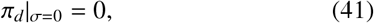

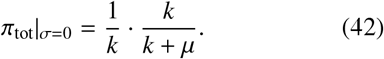

These quantities have natural probabilistic interpretations: two lineages in the same subpopulation of density (that is, the limiting proportion of the total population, as *N* → ∞) *p*, coalesce at rate 2*/p*; each lineage experiences mutations at rate *µ*. They will be identical by descent if and only if there has been no mutation before their eventual coalescence, with probability 2*p*^−1^*/*(2*p*^−1^ + 2*µ*). This gives meaning to *π*_*s*_|_*σ*=0_ when taking *p* = 1*/k*. Individuals from different subpopulations are necessarily non-identical at the neutral marker since subpopulations evolve independently of one another. Thus *π*_*d*_|_*σ*=0_ = 0. The metapopulation probability of identity, *π*_tot_|_*σ*=0_, is given by the probability that the two lineages are from the same subpopulation (with probability 1*/k*), and then coalesce before mutation (with probability *π*_*s*_).

We thus see that in the extreme (if artificial) limit of non-interacting mating type subpopulations, Nei’s gene diversity at the population level (*h*_tot_ = 1 − *π*_tot_) increases as a function of the number of mating types. Meanwhile gene diversity within the mating type subpopulations (*h*_sub_ = 1 − *π*_*s*_) decreases as the number of mating types increases.

#### 3.1.2. Unisexual population

As addressed previously, the probability of identity in a unisexual population is the same as in an asexual population (see Eq. (23)):

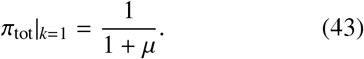

Thus Nei’s gene diversity at the population level is independent of the rate of sexual reproduction in the uni-sexual population. We stress that we do not expect the genetic diversity in general to be independent of the rate of sexual reproduction. Rather, only the diversity at the neutral locus is unchanged, with the issue of genome-wide diversity obscured by Nei’s measure, which focuses on a single locus.

#### 3.1.3 Self-incompatible population

In a facultatively sexually reproducing population with *k* mating types held at constant equal frequency, the probability of identity within the subpopulations and total population are given by Eqs. (20) and (22) respectively. By comparing these quantities to the base case *σ* = 0 (i.e. when *k* asexually reproducing populations held in constant frequency evolve independently of one another, see Eqs. (40-42)), we are led to a natural definition of an effective subpopulation and metapopulation density, as we discuss below.

Assigning a ‘mass’ *N*^−1^ to each individual in the population at the *N*th stage of rescaling, the *density* of a subpopulation *i* is given by *N*_*i*_*/N*. The limiting subpopulation density is given by *p*_*i*_ = lim_*N*→∞_ *N*_*i*_*/N*. In this case of interest, where we artificially hold subpopulations in constant frequency, each subpopulation has limiting density *k*^−1^. The total, or meta-, population density is 1. In the absence of all other interactions, two lineages in an asexually reproducing subpopulation of density *p* are identical by descent with probability *p*^−1^*/*(*p*^−1^ + *µ*). It is reasonable then to say that a population with probability of identity *π* has effective population density

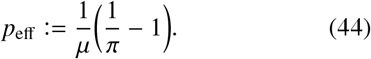

A similar notion of effective population size is defined by Durrett in Section 4.6 of Durrett (2008), where it is called the *heterozygosity effective population size*.

Substituting *π*_*s*_ as given by Eq. (20), the effective sub-population density is

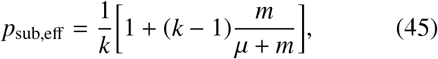

and substituting *π*_tot_ as given by Eq. (22) the effective total population density is

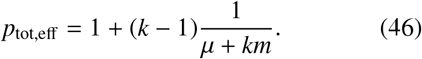

Elegantly, if 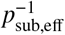 is an integer, the quantities given by Eqs. (20) and (22) correspond to a purely asexually reproducing population divided into 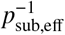 independently evolving subpopulations held in constant frequency with density given by *p*_sub,eff_.

For large *N*, we can translate back to a notion of effective population *size*, writing

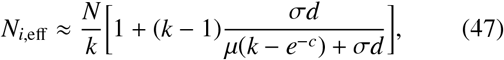

and

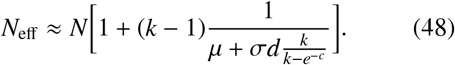

We have used that *m* = *σd/*(*k* − *e*^−*c*^) (see Eq. (19)). *N*_*i*,eff_ is approximately equal to *N/k*, what we would expect for *k* independent asexual populations, multiplied by a factor which is larger than 1, increasing in *k*, and increasing to 1 + *σd/µ*: subpopulations are less susceptible to genetic drift, i.e. invading genotypes are less vulnerable to loss or fixation within subpopulations, when the probability of mutation is small relative to the probability of sex. Similarly, the total effective population size is approximately equal to *N* multiplied by a factor larger than 1; this factor increases approximately linearly in *k*, scaling like *k/*(*µ* + *σd*). We observe that if *σ* = Θ(*k*), i.e. if the rate of sex is on the same order as the number of mating types, then there is no linear growth in the effective population size. Conversely, when the rate of sex is small, the effective total population size can be very large indeed.

In a final remark, we observe that taking *σ*→ ∞, that is, taking the rate of sex to be very large, Eqs. (20) and (22) are both given by the probability of identity for two lineages in a homogeneously mixed asexually reproducing population given by *π*_tot|*σ*=0,*k*=1_. This is consistent with a phenomenon called ‘the collapse of structure’ in structured coalescent models; the reader is directed to the informative summary in Chapter 6 of Etheridge (2011) and the original paper by Nordborg and Krone (2002).

In summary, the Nei’s gene diversity at the population level (*h*_tot_ = 1−*π*_tot_) increases with the number of mating types *k* (for *k ≥* 3, or for all *k* if *σd ≥*2*µ*, i.e. if the rate of recombination is much larger than the rate of mutation), while gene diversity within a subpopulation (*h*_s_ = 1−*π*_s_) decreases as a function of the number of mating types (under the same conditions). However, both these responses are muted by an increasing number of attempted sexual reproductive events per generation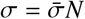.

#### 3.1.4 Comparison of monomorphic populations

In Figures 4 and 5 we compare our analytic results, for total population diversity and subpopulation diversity respectively, to the results of Gillespie simulations. We see excellent agreement across a range of parameters.

**Figure 4:**
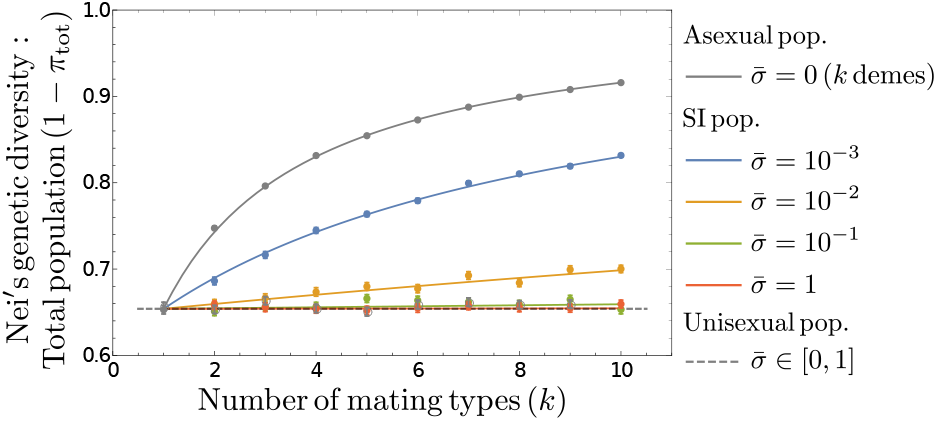
Total population genetic diversity (as measured by Nei’s gene diversity) for asexual, self-incompatible and unisexual populations. Theoretical results (solid lines) well-predict the results of simulations (markers). Theoretical predictions for an asexual population with *k* non-interacting demes are obtained from Eq. (42), for a unisexual population from Eq. (43) and for a population with *k* self-incompatible mating types from Eq. (22). Model parameters: *N* = 7560, 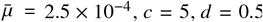. Simulations are averaged over 10^3^ time points sampled regularly over 10^7^ generations following an allowed 10^5^ generation equilibrating period (initial conditions are all individuals homogenous at the neutral marker).

**Figure 5:**
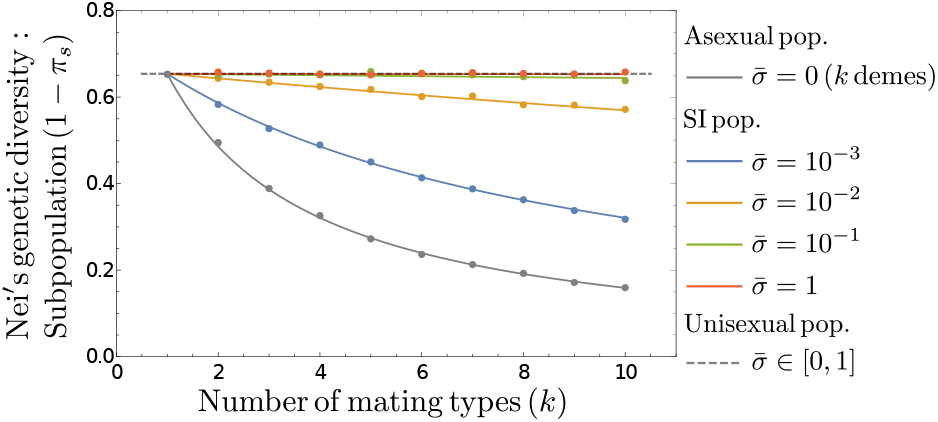
Population genetic diversity in mating type subpopulations (as measured by Nei’s gene diversity) for asexual, SI and unisexual populations. Theoretical results (solid lines) well-predict the results of simulations (markers). Theoretical predictions for an asexual population with *k* non-interacting demes are obtained from Eq. (40), for a unisexual population from Eq. (43) and for a population with *k* SI mating types from Eq. (20). Model parameters are the same as those in Figure 4.

For unisexual populations we find that diversity at the neutral locus is unmodified from the asexual case and thus independent of rate of sexual reproduction. In SI populations, we see that the total population level diversity increases monotonically away from the unisexual baseline as a function of the number of mating types (although at a decreasing rate as more types are added, see Figure 4). While this effect increases as the rate of sexual reproduction decreases, this is entirely a result of the division of the population into mating type subpopulations. (This can be understood intuitively by considering recombination as providing a mechanism for a “migration” event between mating type backgrounds acting as demes.) As a result the genetic diversity in the total population is in fact maximised when sex is entirely absent. However, this is a rather artificial case; in this regime, as we have shown in Section 2.1, there is no selection for mating types and so the mating-type population classes would be lost to drift. Similarly when sexual reproduction is rare and selection for equal mating type frequencies weak, mating types become increasingly susceptible to extinctions as their number increases. Meanwhile in the converse limit of obligate sex, the total population diversity relaxes to the values found in the unisexual and asexual cases. Again this can be intuitively understood by considering “obligate migration” between mating type “demes”, which erodes the effect of population substructure.

While genetic diversity within the total population increases as a function of the number of mating types, the genetic diversity within a mating type subpopulation decreases (see Figure 5). This is a result of smaller sub-population sizes (*N/k*), which simultaneously decreases the effective mutation rate and increases genetic drift within each class. The effect is amplified by low rates of sexual reproduction which increases the genetic isolation of the respective subpopulations.

We note that in both the cases of total population genetic diversity and subpopulation genetic diversity the key parameter with regards to facultative sex is the number of sexual events per generation, 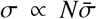, rather than simply the probability a sexual reproductive event is attempted,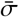.Thus increasing the population size while keeping 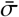 fixed has a similar qualitative effect to increasing 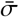 with fixed *N*; the response of *h*_*tot*_ and *h*_*s*_ to increasing *k* is decreased.

### 3.2 Rate of effectual sex

In this section, we seek to understand how the benefits of sexual reproduction experienced may be affected by the presence or absence of mating types and their number. As the benefits of recombination can only be realised when sex occurs between genetically distinct individuals, we calculate the per capita probability that an individual successfully mates with an individual that is genetically distinct at the neutral locus. We term this the probability of effectual sex, which is given by the probability that a compatible mate is successfully located, multiplied by the probability that this mate is not identical by descent at the neutral marker. For each *i* = 1, …, *k*, this probability for an individual with mating type *i* can be expressed as

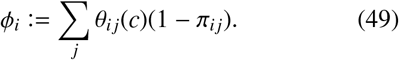

We note that this quantity decreases when the probability of finding a mate decreases (e.g. in SI populations) but increases with increasing genetic diversity between selected mates (caused, for example, by mating type population structure). It is the balance between these competing factors that we will explore in the following subsections. We first explore the behaviour of *ϕ*_*i*_ in the cases of monomorphic unisexual and SI populations, before proceeding to consider how *ϕ*_*i*_ changes in mixed populations of unisexuals and SI mating types.

#### 3.2.1 Unisexual population

Because there is no self-incompatibility in the unisexual population, the probability of finding a mate is one (*θ*_*uu*_(*c*) = 1, see Eq. (25)). Meanwhile the probability of IBD for two randomly selected individuals is *π*_*uu*_ = *π*_tot_ |_*k*=1_, with *π*_tot_| _*k*=1_ given in Eq. (43). Thus the probability of effectual sex in a monomorphic unisexual population is given simply by

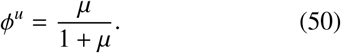

From this we see that *ϕ*^*u*^ is independent of the frequency of sexual reproduction (see Figure 6).

**Figure 6:**
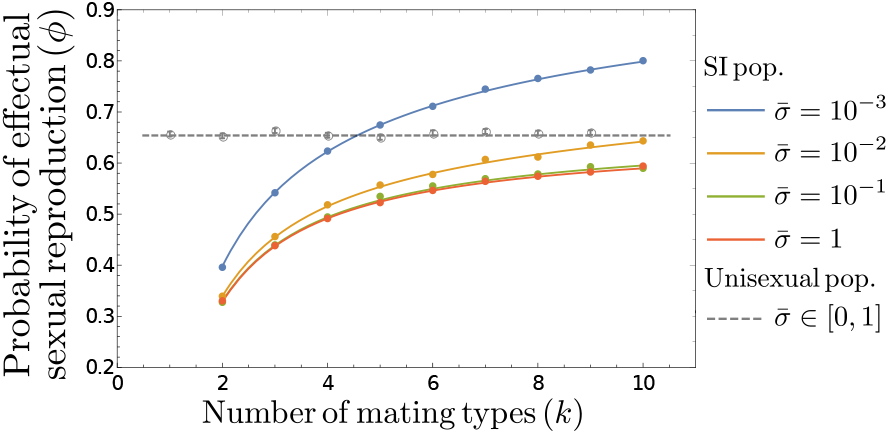
Probability of effectual sexual reproduction, *ϕ*, for monomorphic SI and unisexual populations. Theoretical results (solid lines) well-predict the results of simulations (markers). Theoretical predictions for a unisexual population are taken from Eq. (50) and for a population with *k* SI mating types from Eq. (52). Model and simulation parameters are the same as those in Figure 4.

#### 3.2.2 Self-incompatible population

In the presence of self-incompatibilities, the probability of finding a compatible mate is now generally less than one (see *θ*_*i*_(*c*) in Eq. (17)). Meanwhile the probability of IBD for two individuals selected from distinct and compatible mating type subpopulations *i* ≠ *j* is *π*_*i j*_ = *π*_*d*_, with *π*_*d*_ given in Eq. (21). Thus the probability of effectual sex in a monomorphic SI population is given by

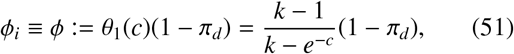

where we note that *ϕ* is the same for all mating types as we have assumed each is held at equal frequency 1*/k* (see Mathematical Analysis). Rewriting this in terms of the original model parameters, we find

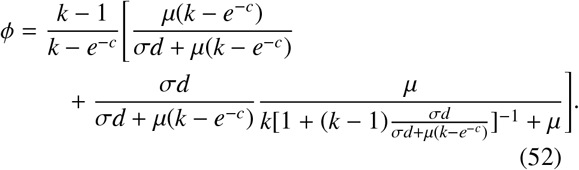

Thus we see that the probability of effectual sexual reproduction increases with decreasing *c* (which increases the probability of locating a compatible mate) and with an increasing number of mating types (which increases the probability of mating with a genetically distinct individual, see Figure 6).

#### 3.2.3 Mixed self-incompatible and unisexual population

In this section we consider a mixed population consisting of *k* − 1 SI mating types and a single unisexual type. As discussed in the Mathematical Analysis section, when calculating the probability of IBD in the mixed population we are forced to consider simplified cases of the model in order to make analytic progress. This is because the unisexual type will always sweep to fixation in the absence of any reproductive benefits to self-incompatibility. We therefore consider three scenarios under which *ϕ*_*i*_ is calculated in turn: a unisexual type held at frequency *α* and *k* − 1 SI mating types each at frequency (1 − *α*)*/*(*k* − 1) in a population at equilibrium; a rare unisexual type arising in a population of *k* − 1 SI types previously at equilibrium; and *k* − 1 rare SI types arising in a population of the unisexual type at equilibrium.

Recall that we denote the probability of IBD as *π*_*d*_ for two individuals sampled from distinct mating type classes, *π*_*s*_ for two individuals sampled from the same mating type class, 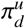 for one individual sampled from a mating type class and another sampled from the unisexual class, and 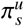 for two individuals sampled from the unisexual class. We then denote by *ϕ* and *ϕ*^*u*^ the probability of effectual sexual reproduction for SI mating types and unisexual types respectively. Using Eq. (49), these probabilities can be expressed as

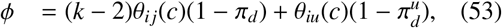

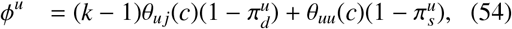

where *i, j* are arbitrary indices representing SI classes (i.e. chosen such that *i* ≠ *j* and *i, j* ≠ *u*) and the forms of *θ*_*i j*_(*c*) and the probabilities of IBD depend on the scenario under consideration.

When the unisexual type is held artificially at a constant frequency and the remaining SI mating types at equal frequencies, expressions for *θ*_*i j*_(*c*) can be taken from Eq. (25) with *p*_*u*_ = *α* and *p*_*i*≠*u*_ = (1 − *α*)*/*(*k* − 1). Substituting these into Eqs. (53-54), we obtain

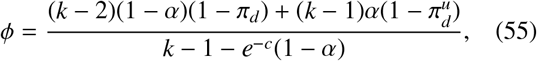

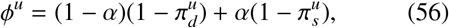

with 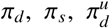 and 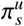 taken as the solution to the Eqs. (I.1-I.4) in Appendix I. The full expressions are too algebraically complex to reproduce here. However, we find the following general patterns. When the frequency of unisexuals is very low (*α* → 0), the rate of effectual sex is greater for unisexuals than SI mating types; *ϕ*^*u*^ > *ϕ* for all *k*. Conversely when the unisexual frequency is very high (*α* → 1), the rate of effectual sex is greater for SI mating types than unisexuals; *ϕ* > *ϕ*^*u*^ for all *k*. The precise value of *α* for which this transition occurs is dependent in a complex manner on the model parameters. We shall explore this more thoroughly in the following subsection.

Next we consider the scenario in which a unisexual mutant emerges in an SI population with *k* − 1 mating types, previously at equilibrium. Expressions for *θ*_*i j*_(*c*) can be taken from Eq. (25) with *p*_*u*_ ≈ 0 and *p*_*i*≠*u*_ = 1*/*(*k* − 1). Substituting these into Eqs. (53-54), we obtain

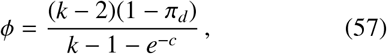

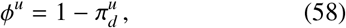

with 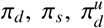 and 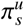 taken from Eqs. (33-36). We find that this qualitatively recapitulates the prediction of Eqs. (55-56) in the limit *α* → 0.

Finally we consider the scenario in which (*k* − 1) mutant SI mating types simultaneously emerge in a resident unisexual population, previously at equilibrium. Expressions for *θ*_*i j*_(*c*) can be taken from Eq. (25) with *p*_*u*_ ≈1 and *p*_*i*≠*u*_ = 0. Substituting these into Eqs. (53-54), along with expressions for 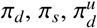 and 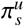 taken from Eqs. (37-38), we obtain

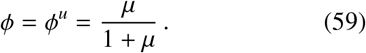

This indicates that upon emergence, each mutant SI type has exactly the same probability of effectual sex as its unisexual ancestor.

We note that while Eq. (59) qualitatively disagrees with the prediction obtained in Eqs. (55-56), (for which *ϕ* > *ϕ*_*u*_ at *α* = 1), this is not unexpected. The approximation in Eqs. (55-56) is obtained assuming that the population is at equilibrium; thus in this limit there is non-zero probability that the SI types (each at infinitely low frequency) carry a distinct neutral genetic marker to those seen in the unisexual population. The approximation in Eqs. (59) on the other hand is obtained assuming that the SI types inherit the neutral marker of their unisexual ancestor; thus in this limit there is zero probability that the SI types (each at infinitely low frequency) carry a distinct neutral genetic marker to those seen in the unisexual population.

#### 3.2.4 Comparison of cases

In Figure 6 we compare our analytic results for the probability of effectual sex, *ϕ*, to the results of Gillespie simulations. Once again, we see excellent agreement across a range of parameters. We see that for SI populations, the probability of effectual sex increases with an increasing number of mating types, *k*, and a decreasing probability of instigating a mating round, 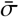, both of which increase the effect of population structure. When the rate of sexual reproduction is extremely low and the number of mating types is moderately high 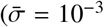, *k* = 5), we see that the probability of effectual sex in the SI population can in fact exceed that of unisexual populations. Thus while the unisexual population engages in more sexual reproduction overall, many of these opportunities to realise the benefits of sex are essentially wasted by same-clone mating.

In Figure 6, *N* and *c* are held fixed. In fact *c* = 5 in these plots, which is approximately equivalent to mass-action encounter rates (a large cost to mate finding). In Figure 7 we explore how the relative advantage to the SI populations over the unisexual population in terms of the probability of effectual sex *ϕ* varies with these parameters. We note that since the key parameter in Eqs. (50) and (52) is in fact the *number* of instigated mating rounds per generation, 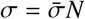, for the SI population to have a higher *ϕ* than the unisexual population for *k* = 2 requires either extremely low 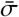 or relatively low *N* (see Figure 7(A-B)). However, an increased number of mating types can appreciably alleviate this requirement (see Figure 7(C-D)).

**Figure 7:**
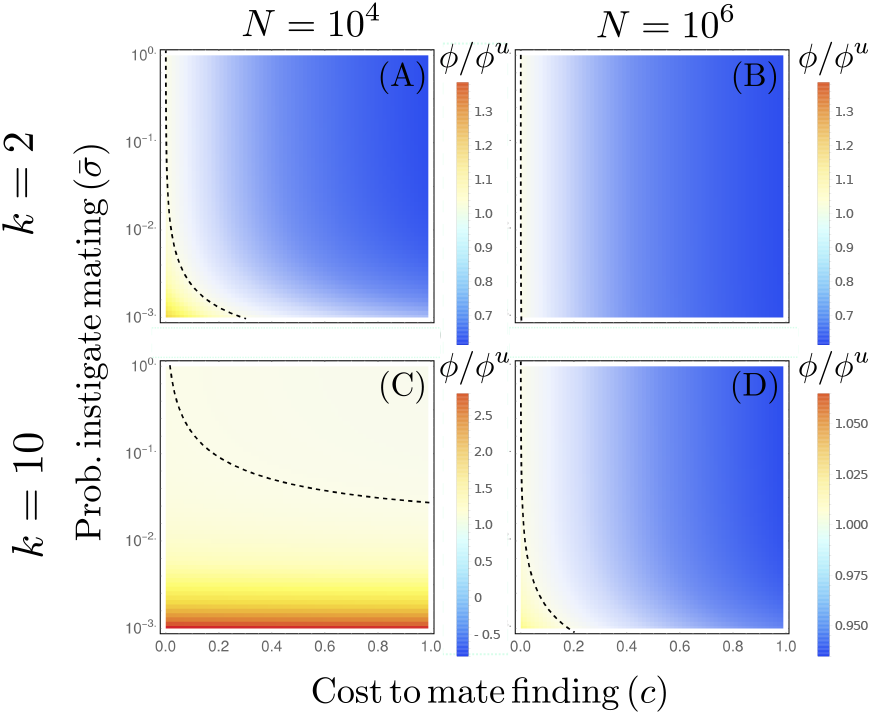
Theoretical predictions for the ratio of probabilities of effectual sexual reproduction for SI mating types, *ϕ*, over unisexual types *ϕ*^*u*^, in monomorphic populations (see Eqs. (50) and (52)). In all panels 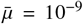 and *d* = 1*/*2. The ratio *ϕ/ϕ*^*u*^ decreases with increasing population size (compare (A) to (B) and (C) to (D)), but increases with a greater number of mating types (compare (A) to (C) and (B) to (D)).

In the above paragraphs, we have started to identify the potential advantages garnered by self-incompatibility over unisexuality at the population level. However, we have done so by comparing populations consisting of purely SI or unisexual types. We next discuss how these insights are altered in the mixed population.

As previously described, we began by calculating the probability of effectual sex (see Eqs. (55) and (56)) for unisexual and SI types (*ϕ*^*u*^ and *ϕ* respectively), under the condition that the frequencies of each of the types were fixed. This is equivalent to making a *quasi-static approximation* that assumes that the population reaches an equilibrium in the distribution of neutral markers on a much faster timescale than that on which the frequency of unisexuals changes. In biological terms this can be understood as assuming very weak selection for unisexuals over SI mating types, or vice versa.

The results of this quasi-static approximation are illustrated in Figure 8, Panels (b-c). The probability of effectual sexual reproduction for SI types (solid lines) behaves in a qualitatively similar manner to that in the case of a monomorphic SI population (see Figure 6); *ϕ* increases with an increasing number of mating types *k* and a decreasing probability of instigating a mating round 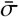. However, we see qualitatively different behaviour of *ϕ*^*u*^ when comparing the monomorphic and mixed population cases. When rare, unisexual types are now able to exploit the diversity generated by the SI types (see Figure 8, Panel (b), dashed lines) to increase *ϕ*^*u*^ beyond what is observed in a monomorphic unisexual population (see Figure 6). As, in addition, they are compatible with the entire population, their rate of effectual sex can exceed that of the SI types. However, as the unisexual frequency increases (see Figure 8, Panel (c)) their probability of mating with genetically similar individuals increases, while simultaneously the costs paid by SI types in terms of restricted mating opportunities decreases. Together these factors can combine to give a higher rate of effectual sex for the SI population than the unisexual population when mating types are numerous, and sex and the SI types themselves are rare.

**Figure 8:**
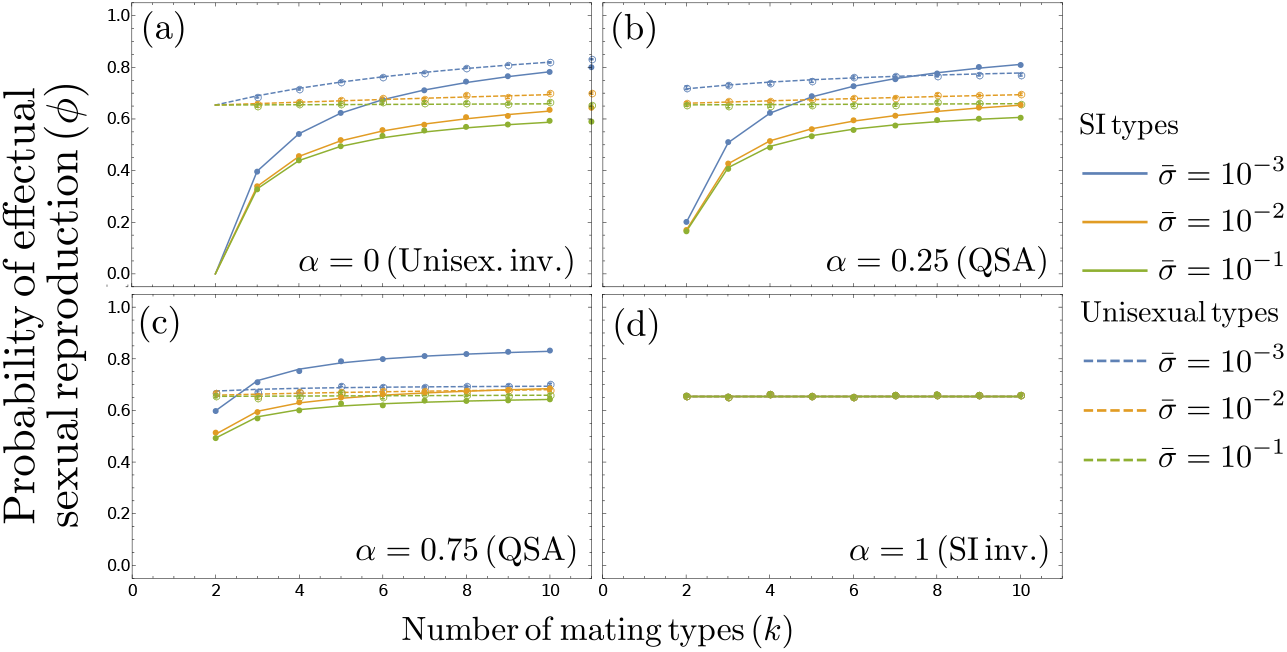
Probabilities of effective sexual reproduction, *ϕ* and *ϕ*^*u*^, for mixed populations with a unisexual type at frequency *α* and (*k* − 1) SI mating types at frequencies (1 − *α*)*/*(*k* − 1). Theoretical results (solid lines) well-predict the results of simulations (markers). Panel a: Scenario of a rare unisexual type invading resident population of mating types, with theoretical results taken from Eqs. (57-58). Panels b-c: Scenario of each type held at a constant frequency so that the population has time to equilibriate to a quasi-stationary state, with theoretical results taken from Eqs. (55-56). Panel d: Scenario of a set of rare SI types invading a resident unisexual population, with theoretical results taken from Eq. (59). Model and simulation parameters are the same as those in Figure 4.

While the quasi-static approximation described above assumes that the population is at equilibrium, we also developed out of equilibrium expressions for the probability of effectual sex, *ϕ*^*u*^ and *ϕ*, under the scenarios of a unisexual type invading a resident population of SI mating types (see Eqs. (57-58)) and of SI mating types invading a unisexual population (see Eq. (59)). These results are illustrated in Figure 8, Panels (a) and (d) respectively. We see that a unisexual invader is able to exploit the genetic diversity generated by resident SI types (see Panel (a)), in qualitative agreement with the quasistatic approximation. However, invading mating types initially receive no advantage in terms of an increased *ϕ* (see Panel (d)), in disagreement with the quasi-static approximation. This implies that while SI types may experience a higher rate of effectual sex than the unisexual population when the SI types themselves are rare, during their invasion this advantage is contingent on them surviving long enough to genetically differentiate from their unisexual ancestors.

In summary we can paint the following picture. When SI mating types first emerge in a unisexual population, they will have an equal rate of effectual sex to the resident population (see Figure 8, panel (d)). As they are rare they pay no costs in terms of reduced mating opportunities, but simultaneously receive no benefits from self-incompatibility, having not had time to differentiate themselves from their unisexual ancestors. Any initial growth in their frequencies will be driven primarily by drift. However, if these mating types can survive at low frequencies for a sufficient amount of time, they will differentiate genetically from their unisexual ancestors. In this case when the cost to mate finding is low and sex is rare 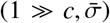, SI types will mate more frequently with genetically distinct individuals than unisexuals (see Figure 8, panel (c)). If the benefits of sex lie in recombination, then the mating types may experience a selective advantage over their unisexual ancestors. Although this may temporarily drive an increase in the frequency of SI types, this advantage is short-lived. As the frequency of SI types increases, their opportunities to find a compatible mate decrease. Simultaneously the unisexual type increasingly exploits the genetic diversity generated by the SI types. Together these factors can drive the probability of effectual sex in unisexuals above that of the invading mating types (see Figure 8, panel (b)), an effect that becomes ever stronger as the SI mating types approach fixation.

Conversely, when unisexual types first emerge in a population of SI mating types, they are able to exploit the genetic diversity of the resident population to obtain all of the benefits of self-incompatibility (decreased probability of clone-mating) while not paying the costs of missed mating opportunities (see Figure 8, panel (a)). They will thus receive a selective advantage and increase in frequency. However, as they increase, their probability of effectual sex drops below that of their SI ancestors, an effect that becomes stronger as the unisexual types approach fixation (see Figure 8, panel (b) to (c)). If the benefits of recombination are sufficiently strong, this will generate selection against the unisexual type. Taken together, these results suggest that we may in fact see negative frequency dependent selection for self-incompatibility in haploids.

## 4. Discussion

The existence of sexual self-incompatibility between gametes has posed an evolutionary puzzle for many years, and a variety of theories to explain the evolution of this discriminatory behaviour have been proposed (Billiard et al., 2011). Among these, the hypothesis that mating types, by preventing haploid selfing, enhance the advantages of recombination has remained poorly investigated theoretically (Billiard et al., 2012). With the precise benefits of recombination still a widely debated topic, the reason for this deficit is clear; a full investigation of the theory in fact requires distinct studies testing it within the context of each separate hypothesis for the evolution of recombination. In this paper we have circumvented this issue by first studying a more general problem; as stated succinctly in Billiard et al. (2012) *“if recombination has an advantage, whatever it is, individuals gain this advantage only when the haploid genomes that are recombining are not strictly identical”*. We have shown that although self-incompatibility decreases the probability of successful sexual reproduction, it can in fact increase the probability of effectual sexual reproduction between non-clone mates, even exceeding the rates of effectual sexual reproduction in unisexuals. However, the strength of this effect is highly dependent on population parameters including its effective size, the rate of sexual reproduction and the population composition.

We have demonstrated that monomorphic unisexual populations are less diverse than populations composed of SI mating types and further, that the probability of effectual sexual reproduction with an individual genetically distinct at a neutral marker can be lower in a unisexual population than in an SI population. Both these population-level benefits to self-incompatibility vary in a qualitatively similar way as the remaining parameters in the model are varied. Increasing the number of mating types and decreasing the number of sexual events per generation 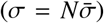 increases the relative benefits to self-incompatibility over unisexuality. Meanwhile, decreasing the cost to mate finding (i.e. allowing more opportunities for the SI types to find a compatible mate) has a particularly prominent effect on the probability of effectual sex in the SI population, allowing it to more readily exceed that in a unisexual population.

The above discussion is limited to considering population level benefits of self-incompatibility over unisexuality. However, for an evolutionary perspective, we must instead understand the costs and benefits of SI at an individual level. Here our simplified “recombination hypothesis free” approach (where the benefits of recombination have not been explicitly defined) runs into some trouble; as we have shown, without any emergent benefits to mating with genetically distinct individuals, the unisexual type is positively selected for. Although the strength of selection against self-incompatibility is decreased with lowered costs to mate finding, the unisexual type will always sweep to fixation in finite time.

To make progress, we investigated how the rate of effectual sex (an individual benefit) varied between unisexual and SI types in a population held artificially at a polymorphic equilibrium. This approximation to the dynamics of the full model (in which the frequencies of unisexuals and SI mating types are allowed to vary) is likely to hold when selection for either unisexual or SI types is relatively weak. We found that in this mixed population, the unisexual individuals can exploit the diversity generated by mating types, obtaining a higher rate of effectual sex than in their equivalent monomorphic populations. This scenario is reminiscent of the ‘tragedy of the commons’, a comparison previously utilised to explain selection for low-recombination rate modifiers in general models for the evolution of recombination (Kokko, 2020). While this leads to unisexuals having a higher probability of effectual sex than their SI mating type competitors when rare, this advantage is frequency dependent; as the frequency of unisexuals increases, their rate of effectual sex dips back below that of their SI competitors. While the precise frequency at which this transition occurs depends in a complex way on the model parameters, our intuition from the case of monomorphic populations carries over: the advantage of self-incompatibility over unisexuality diminishes as the number of mating types decreases, and the number of sexual events per generation 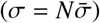 and costs to mating finding (*c*) increase.

We next turn our attention to the key biological motivation for this paper - in the context of the sex-advantage enhancer hypothesis: (i) how did SI first evolve and (ii) how does SI remain robust against invasion?

Regarding the first question, we note that initially (when rare) a mutant SI type would only be weakly selected against. However, in order to drift to more substantial frequencies the costs to SI of mate finding would have to be low. This requires many mating rounds and thus suggests that self-incompatibility first evolved in an aquatic environment, in which individuals have more opportunities to actively find a compatible mate. Such an environment is the natural case for algae. While fungi are more often associated with terrestrial environments, empirical evidence suggests that early fungal ancestors lived in water (Naranjo-Ortiz and Gabaldón, 2019). With the MAT locus that determines mating types in fungi appearing ancient and displaying a remarkable conservation of core homeodomain proteins over trans-specific evolutionary periods (Fraser et al., 2004), it is conceivable that self-incompatibility in fungi arose in just such an environment. However, in order to see selection for self-incompatibility, it is not sufficient to simply have low costs; self-incompatibility must also have benefits.

As we have addressed, if the SI mating types can be maintained in the population for a sufficient amount of time at low frequencies, they can obtain a higher rate of effectual sex compared to their unisexual ancestors. If genetic recombination with non-clone mates is sufficiently advantageous this could generate selection for mating types. However, the magnitude of this effect is strongly dependent on the remaining population parameters. Crucially this benefit to self-incompatibility requires that the number of sexual events per generation,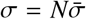, is low. While it seems natural to assume that when self-incompatibility first evolved, the rate at which sexual reproduction and mate finding was triggered was very low 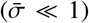, low effective population sizes (e.g. *N* ≪ 10^6^, see Figure 7) are harder to justify biologically, especially if we consider these dynamics occurring in a well-mixed aqueous environment. Such low population size values are more consistent with a spatially structured population, within which the costs to mate finding are expected to be larger. We proceed with this caveat in mind.

Assuming that the SI mating types are selected for at this stage of their invasion trajectory, we emphasise that this advantage over unisexuals is necessarily fleeting. Once the SI types reach a critical frequency, unisexual types will be able to parasitise their diversity and regain the upper hand. Thus in terms of self-incompatibility having a higher probability of displacing unisexuality than vice-versa, the very best that can be hoped for is that this transition occurs at a high frequency of SI types, far into their invasion trajectory. We note that once a resident population of two mating types has established, an evolutionary pressure for the number of mating types to increase arises. New mating types experience a selective advantage both in terms of increased mating opportunities relative to the two residents and a (shared) increase in the probability of mating with a genetically distinct individual. There is then selection for an ever greater number of mating types, which are only kept bounded by genetic drift (Constable and Kokko, 2018; Czuppon and Rogers, 2019).

We now consider the second question, of how self-incompatibility can remain robust against invasion from unisexual, or homothalic, mutants. Our model suggests that in fact self-incompatibility may never be truly robust against invasions from unisexuals, as unisexuals can always free-ride on the genetic differentiation of resident SI mating types. However, fixation of unisexuals might be prevented if their probability of effectual sexual reproduction drops below that of SI residents at sufficiently low frequency. We have shown that in this context, the presence of multiple resident mating types can markedly enhance this possibility, such that SI types can maintain a higher rate of effectual sex even as the number of instigated sexual events per generation becomes moderately high (see Figure 7). It is therefore interesting to note that among the Basidiomycota, which are characterised not only by their comparatively high rates of sexual reproduction (Lee et al., 2010), but also by their large number of mating types (Coelho et al., 2017), homothalism is relatively rare (James, 2007). Homothalism is also far from ubiquitous in isogamous algae, with only a handful of species known to employ this indiscriminate mating strategy (Sekimoto et al., 2012). Here our model suggests qualitatively that self-incompatibility may be protected by the rarity of sex in these species (Hasan and Ness, 2020). However, in a quantitative sense, our model predicts that this rarity should be offset by the large effective population sizes observed in isogamous algae (Ness et al., 2012).

Taken together, we see from the above discussion that the sex advantage enhancer hypothesis may agree qualitatively with observed scenarios under which self-incompatibility first evolved (i.e. an aquatic environment with low costs to mate searching and very rare sexual reproduction), is maintained (i.e. in populations with a low number of sexual reproductive events per generation and a large number of mating types) or is displaced by homothalism (i.e. populations with more frequent sexual reproduction, a low number of resident mating types and high costs to mate finding). Although in a quantitative sense we see that our model predicts that self-incompatibility may be selected for over a rather restricted range of biologically reasonable parameters, we have made a number of simplifications in order to make the problem tractable, which may provide an overly stringent estimation of when self-incompatibility may be selected for. Below we detail some of these key simplifications, as well as discussing their potential effects.

First, for simplicity, we have considered only the dynamics of a single neutral genetic marker, and defined sex as ‘ineffectual’ when two mating individuals carry the same marker. In reality, particularly in populations undergoing recombination, it is of course possible that while individuals are identical at the neutral marker, they are genetically distinct throughout the rest of their genome. Thus our estimate very much provides a lower bound for the rate of potentially beneficial sex between non-clone mates. Despite this we believe that the qualitative intuition that we have developed will remain valid; frequent sexual reproduction erodes population structure, and is thus likely to diminish the advantage to self-incompatibility in well-mixed populations.

Second, while we have assumed a well-mixed population, it is clear that spatial structure, by increasing the probability of mating with genetically related individuals, is likely to be a strong driver of the evolution of self-incompatibility. Within our modelling framework we view this effect as being equivalent to small population sizes. While it would be interesting to consider how explicitly accounting for spatial structure would affect the results we have described here, we note that these additional insights would likely come at the cost of generality, with any model necessarily having to account for species-specific details of asexual growth environment, mate searching and non-local dispersal.

Third, we note that our modelling approach is based on the assumption that mating with a genetically distinct individual increases the advantages of sexual reproduction. However, the benefit of sexual reproduction is not restricted to such recombination advantages. In many species, stress-induced sexual reproduction has evolved to be coincident with an environmentally resistant dormant form. In cases such as these, the primary advantage to sex (survival) is achieved regardless of the genetic identity of the partner. For instance, in the yeasts *Saccharomyces cerevisiae* and *S. pombe*, sexual reproduction is followed by the formation of environmentally resistant spores (Gerber and Kokko, 2018). Thus while these species fall well within the range where we would expect self-incompatibility to be favourable (with rare rates of sexual reproduction (Tsai et al., 2008) and spatially structured populations with small local effective population sizes (Nieuwenhuis et al., 2018)), mating type switching has still evolved in these species (Nieuwenhuis and Immler, 2016). This homothalism, while metabolically costly during asexual reproduction, provides reproductive assurance and survival over the longer term and can thus still be selected for (Nieuwenhuis et al., 2018).

Finally, and perhaps most importantly, our model does not include any explicit benefit to genetic recombination. This was very much a conscious decision, as it allows us to avoid ‘baking in’ an advantage to non-self mating, in contrast to earlier studies (Nauta and Hoekstra, 1992; Czaran and Hoekstra, 2004). While our approach falls short of what must be the ultimate goal of investigating how self-incompatibility can be selected for in models in which the benefits of recombination are emergent (i.e. arise under a recombination hypothesis framework), our results nevertheless provide intuitive insight as to how these models might behave. For instance, consider a prototypical model of the Red Queen Hypothesis in a haploid population, with 2 triallelic loci and with selection for recombination generated by matching-allele interactions (Lively, 2010). If sex is regular, the frequencies of the genes in each mating type subpopulation will be roughly equal. In this scenario we might expect there to be no advantage to self-incompatibility, only a cost in terms of mating opportunities. However, if sex were rare, drift would drive deviations in genotype frequencies between mating type subpopulations, increasing the probability of mating with a genetically distinct individual under self-incompatibility.

Our hope is these challenges, such as accounting for multiple genetic markers and benefits of recombination, will be tackled in the near future. In particular we hope that the results of this paper will help guide research that specifically looks at the evolution of SI within each framework for the evolution of recombination. Of key importance will be the incorporation of facultative sex into recent developments in our understanding of how mating systems affect the spread of beneficial mutations in obligately sexual species (Zhang et al., 2020). With further theoretical advancements in our understanding of facultative sex now coming to the fore (Hartfield et al., 2018), we are positive about the potential for progress in this field.

## Acknowledgements

All authors would like to thank the Probability Meets Biology workshop at the University of Bath, where this project was initially formulated. GWAC thanks the Leverhulme Trust for funding through the Leverhulme Early Career Fellowship. IL acknowledges ANID/Doctorado en el extranjero doctoral scholarship, grant number 2018-72190055.

## Code and data availability

All code and data used to generate the figures in this paper can be found in the following repository: https://github.com/gwaconstable/GeneticDiversityMT.git

## Appendix A. Derivation of the deterministic dynamics of self-incompatibility alleles

Initially we consider only the dynamics of the mating type frequencies (and not those of the neutral marker) and we can therefore neglect mutation and recombination. This leaves us only concerned with the reproductive events that change the mating type frequencies. We conduct the following analysis using Model A (see Figure 1), where we assume that the progeny of reproductive events can replace any individual in the population.

Recall that each of the *N* individuals reproduces asexually at rate 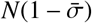, meaning that an asexual reproduction event occurs in the population at total rate 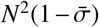. When such an event occurs, an individual of type *i* is picked from the population with a probability proportional to its frequency, *N*_*i*_*/N*, while simultaneously another individual of type *j* is picked to die. Thus the probability per unit time of a type *i* increasing by one and type *j* decreasing by one from asexual reproduction is simply

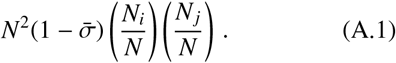

Meanwhile, each of the *N* individuals attempts to reproduce sexually at rate 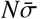, meaning that a sexual reproduction event occurs in the population at total rate 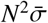. We first consider the case of *k* SI mating types.

When a sexual reproduction event occurs, an individual of mating type *i* is picked from the population with a probability proportional to its frequency, *N*_*i*_*/N*. The probability that it successfully finds a compatible partner of type *j* ≠ *i* is then given by 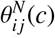 (see Eq. (2)). The offspring of this union inherits the mating type of parent *i* with a probability 1*/*2. This progeny displaces an individual of mating type *l* with a probability *N*_*l*_*/N*. Of course mating type *i* can also increase if instead mating type *j* ≠ *i* is chosen for sexual reproduction (with probability *N*_*j*_*/N*) and locates a type *i* as a mate (with probability 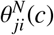, with the progeny again inheriting the mating type of parent *i* with probability 1*/*2. Summing over all potential partner types *j* ≠ *i*, the probability per unit time of mating type *i* increasing by one and mating type *l* decreasing by one from sexual reproduction is given by

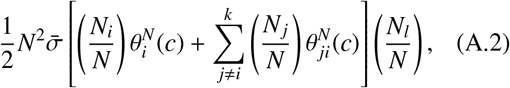

where 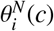 is given in Eq. (3).

Taking the sum of the above expressions, we can now write the probability per unit time that the system transitions from state *N* = (*N*_1_, *N*_2_, …, *N*_k−1_) to *N*^(*i, j*)^ = (…, *N*_i_ + 1, …, *N*_*j*_ − 1, …) as *T*_*i j*_(*N*), where

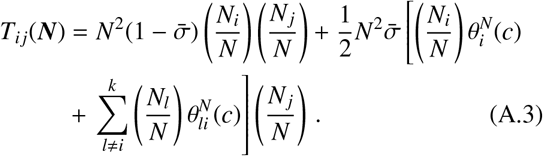

The probability of being in a state ***N*** at time *t*, Φ_***N***_(*t*), is then governed by the master equation:

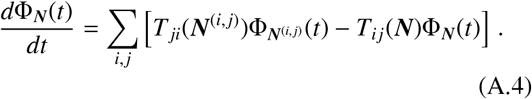

We now make a transformation into *τ* = *Nt* and the approximately continuous set of variables *p*_*i*_ = *N*_*i*_*/N*, such that transitions now occur between neighbouring states ***p*** = (*p*_1_, *p*_2_, …, *p*_*k*−1_) and ***p′*** = (…, *p*_*i*_ + 1*/N*, …, *p*_*j*_ − 1*/N*, …). Conducting a Taylor expansion of the master equation (A.4) in 1*/N* and truncating after leading order (McKane et al., 2014), we obtain

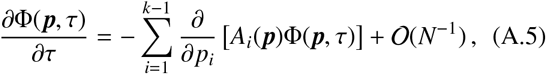

Where A_*i*_(*p*) is given by

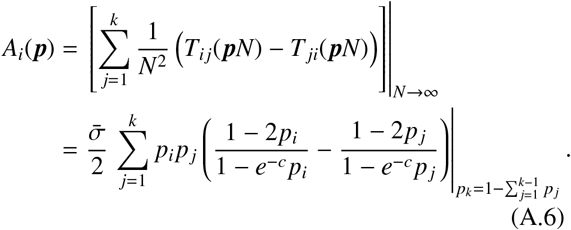

Eq. (A.5) is equivalent to the ODE (McKane et al., 2014)

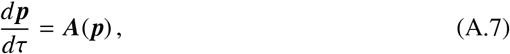

in agreement with Eq. (4) in the main text.

The system Eq. (A.7) has a fixed point at ***p*** = ***p***\* with 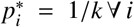 Note that as *c*→ 0 (i.e. each mating type finds a compatible sexual partner with certainty) a reproductive advantage for rare mating types is still maintained. This is because although an individual of a rare type is as likely to find a partner as the common types 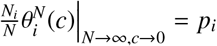 in Eq. (A.3)), a rare type is in fact more likely to be found as a partner than its competitors 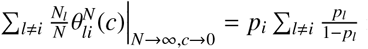 in Eq. (A.3)).

We next calculate the Jacobian of ***A***(***x***) evaluated at the fixed point 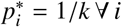. We obtain the (*k* − 1) × (*k* − 1) diagonal Jacobian *J* with

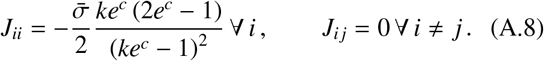

Being diagonal, we can simply read off the (*k* − 1) eigenvalues of *J* as *λ*_*i*_ = *J*_*ii*_.

We next calculate the dynamics in the case of *k* − 1 SI mating types, and a single *k*^*th*^ unisexual type. We can proceed in analogous manner to the previous case, however with modified transition rates that take account of the fact that the unisexual type *k* finds a compatible partner with probability one during a mating round. We Find

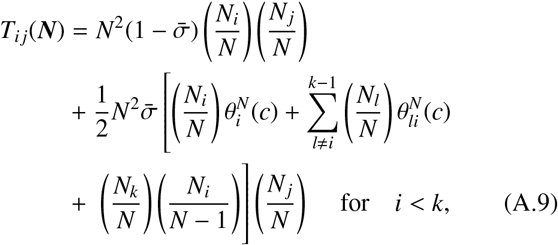

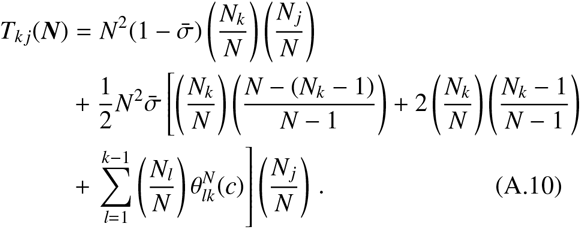

Following the same steps as before (that is, expanding the master equation (A.4) in 1*/N* and truncating after leading order) we now obtain a modified form for *A*_*i*_(***x***) that accounts for the dynamics of the unisexual type:

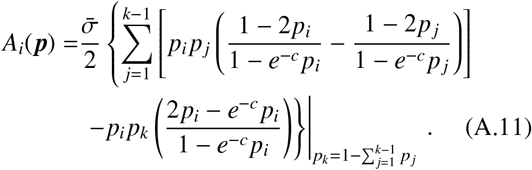

This system features two fixed points of biological interest.

The first fixed point corresponds to the coexistence of the *k* –1 SI types (*p*_*i*_ = 1*/*(*k* -1) for all *i < k, p*_*k*_ = 0). Evaluating the system’s Jacobian at this fixed point we find the eigenvalues

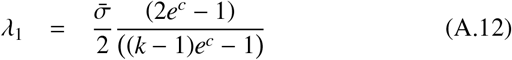

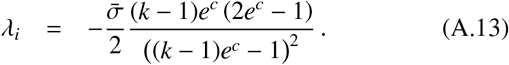

The first eigenvalue is always positive, indicating that the *k* − 1 coexisting SI types are always susceptible to invasion by the unisexual type, which sweeps to fixation.

The second fixed point corresponds to the fixation of the unisexual type (*p*_*i*_ = 0 for all *i < k, p*_*k*_ = 1). This fixed point is only stable at second order in a Taylor expansion about the fixed point, as when extremely rare self-incompatibility carries little cost with respect to missed mating opportunities.

## Appendix B. Diffusion processes

A *k*-dimensional diffusion process {*X*(*t*) = (*X*_1_(*t*), …, *X*_*k*_(*t*))} _*t*≥0_ is a continuous time Markov process with continuous sample paths. It is described by two groups of quantities, called the drift and diffusion coefficients, which describe the mean and covariance in the change in each coordinate *X*_*i*_(*t*) over an infinitesimal time period: if for each *i, j* = 1, …, *k* and *h >* 0 one sets Δ_*h*_*X*_*i*_(*t*):= *X*_*i*_(*t* + *h*) − *X*_*i*_(*t*), and if

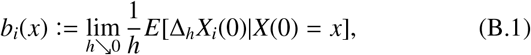

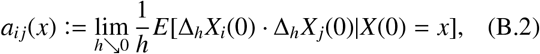

then {*X*(*t*)}_*t*≥0_ is described as a solution to the system of stochastic differential equations

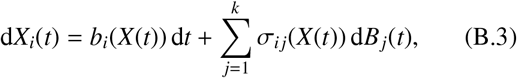

where *B*_1_, …, *B*_*k*_ are independent Brownian motions and the coefficients *σ*_*i j*_, *i, j* = 1, …, *k*, are defined by the relation *a*(*x*) = *σ*(*x*)*σ*(*x*)^*T*^. The coefficients *b*(*x*) = (*b*_*i*_(*x*))_*i*=1,…,*k*_ and *a*(*x*) = (*a*_*i j*_(*x*))_*i, j*=1,…,*k*_ are referred to as the drift and diffusion coefficients respectively. Approximately speaking, we may interpret this as saying that

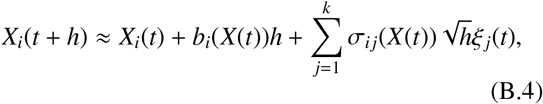

where for each *j* = 1, …, *k* and *t >* 0, *ξ* _*j*_(*t*) is a Gaussian random variable with mean zero and unit variance such that *ξ*_*i*_(*t*) and *ξ* _*j*_(*s*) are independent if either *i* ≠ *j* or *t* ≠ *s*. The diffusion may equally be characterised by its infinitesimal generator *L* defined by

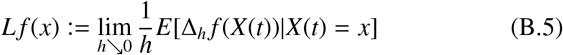

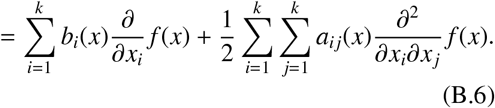

The reader is referred to Karlin and Taylor (1981) and Durrett (1996) for a further introduction to diffusion processes.

Diffusion processes often arise as the limit of a sequence of Markov processes with increasingly frequent but increasingly small jumps; the canonical example is Brownian motion, which arises as the limit of a suitably rescaled simple random walk. Suppose that we have a sequence of continuous time Markov processes 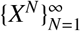, where, for each *N*, the process *X*^*N*^ takes values in *S* _*N*_ ⊆ ℝ^*k*^. If we define for *x* ∈ *S* _*N*_, *i, j* = 1, …, *k*,

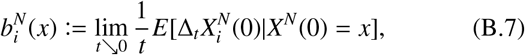

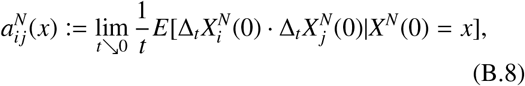

and suppose that for all *R <* ∞,

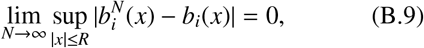

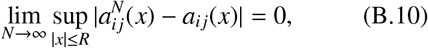

and

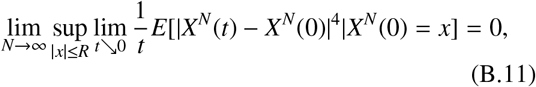

then the sequence of processes *X*^*N*^ approximates the solution *X* to the system of equations (B.3). This is a version of the seminal result of Stroock and Varadhan, see e.g. Durrett (1996), and a full statement is given in the next appendix. The fourth moment condition can be relaxed, although, as in our case, it is often easy to check.

## Appendix C. The Stroock–Varadhan Theorem

In this appendix, we give a precise statement of a version of the Stroock–Varadhan Theorem for continuous time Markov processes; it follows after combining the results of Theorem 7.1 and Lemma 8.2 in Durrett (1996), Chapter 8. The Stroock–Varadhan Theorem provides the conditions under which a sequence of Markov chains, in either discrete or continuous time, converge to a solution to a martingale problem; a further result states when such a solution may be expressed as a solution to a system of stochastic differential equations. An analogous version of the result we state exists for discrete time processes; the reader is directed to Chapter 8 of Durrett (1996) for further details.

Let 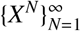 be a sequence of continuous time Markov processes, where for each *N* the process *X*^*N*^ takes values in [0, 1]^*k*^ ⊆ ℝ ^*k*^. Define for each *N* = 1, 2, …, *x* ∈ [0, 1]^*k*^, *A* ⊆ ℝ ^*k*^, *x ∉ A* a sequence of measures

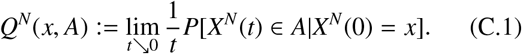

The measure *Q*^*N*^ (*x, A*) records the instantaneous rate at which the process *X*^*N*^ jumps from a state *x* into the subset

A. Let us further define the sequence of moments

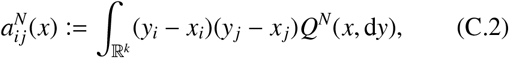

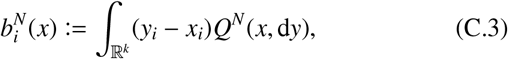

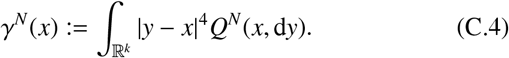

Let us suppose that

A. for each *N*,

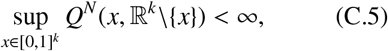
B. *b*_*i*_ and *a*_*i j*_ are continuous coefficients for which, for every initial condition *x* ∈ [0, 1]^*k*^, there exists at most one continuous process *X* on [0, 1]^*k*^ such that *X*(0) = *x* and

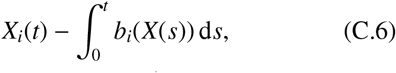

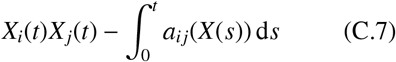

are both local martingales for each *i, j* = 1, …, *k*, and for each *i, j* = 1, …, *k*, it holds that
C. 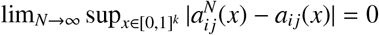 and 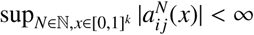;,
D. ((D) 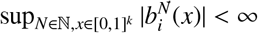 and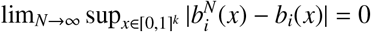
E. 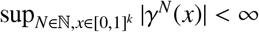 and 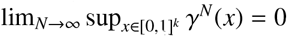

If, in addition to conditions (A-E), it holds that *X*^*N*^ (0) = *x*^*N*^ → *x* in ℝ^*k*^ as *N* → ∞ then, in the same limit, *X*^*N*^ ⇒ *X*, where *X* is the unique continuous process such that *X*(0) = *x*, and (C.6) and (C.7) are local martingales for each *i, j*. Here, ‘ ‘ denotes weak convergence in the space of right-continuous paths *ω*: [0, *T*] → ℝ^*k*^ with left limits (so-called càdlàg paths) equipped with the Skorokhod topology. The space is frequently denoted *D*([0, *T*], ℝ^*k*^) and the reader is directed to Section 8.6 of Durrett (1996), Chapter 8, for further details of this notion of convergence.

Let us now address the circumstances under which condition (B) is satisfied. Defining *σ* such that *a* = *σσ*^*T*^, Theorem 4.5 of Durrett (1996), Chapter 5 tells us that there is a one-to-one correspondence between processes *X* with *X*(0) = *x* satisfying the condition that (C.6) and (C.7) are local martingales, and weak solutions to the system of stochastic differential equations defined in Eq. (B.3). Uniqueness in distribution of solutions to Eq. (B.3), then, guarantees uniqueness of such a process *X*. Verifying (B) is thus equivalent to verifying that

(B’) *b*_*i*_ and *a*_*i j*_ are continuous coefficients for which, for every initial condition *x* ∈ [0, 1]^*k*^, solutions to the system of Eqs. (B.3) with *a* = *σσ*^*T*^ and *X*(0) = *x* taking values in [0, 1]^*k*^ are unique in distribution.

One approach to proving such uniqueness is the method of duality, outlined in the following appendix.

## Appendix D. The method of duality

In this appendix, we outline a working version of the method of duality. This technique, which may initially appear contrived, has found widespread applications in proving uniqueness in distribution of solutions to stochastic differential equations and, more generally, martingale problems. Here, we state a special case of a more general version appearing in Section 4.4 of (Ethier and Kurtz, 1986).

Uniqueness in distribution of solutions to this system of equations is implied by uniqueness of one-dimensional distributions (see, for example, Theorem 4.2 of Ethier and Kurtz (1986), Chapter 4); that is, if 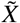 and are two solutions to the system of Eqs. (B.3), it is sufficient to show that, for each *t* ≥ 0,

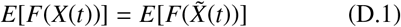

for a sufficiently large class of functions *F*. For example, the set of all smooth and compactly supported *F*: ℝ^*k*^ → ℝ would be one such class (although much smaller classes do exist, as we will make use of later).

Fix *x* ℝ^*k*^. We outline a method for proving uniqueness in distribution of solutions to the system of Eqs. (B.3) with initial condition *X*(0) = *x*. Let *f*: ℝ^*k*^ × ℝ^*l*^ → ℝ and *β*: ℝ ^*l*^ → ℝ be functions. Take *y* ∈ ℝ^*l*^ and suppose that

(F) there exists an ℝ^*l*^ -valued Markov process *Y* such that *Y*(0) = *y* and, for every [0, 1]^*k*^-valued solution *X* to the system of Eqs. (B.3) with initial condition *X*(0) = *x, X* is independent of *Y* and

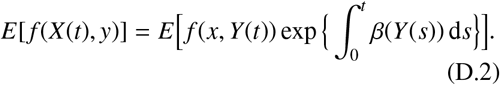

If, furthermore, for each *t* ≥ 0,

(G) 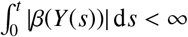 almost surely,

(H) *E*[|*f* (*X*(*t*); *y*)|] > ∞, and and

(I) 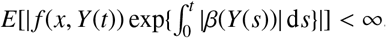

then we say that solutions to (B.3) with initial condition *X*(0) = *x* are dual to the process *Y* with respect to the functions *f* and *β*. Thus the problem of proving uniqueness of solutions to (B.3) with initial condition *X*(0) = *x* is transformed into one of establishing the existence of *Y* for a large enough choice of initial conditions *y*: if such a *Y* exists for all initial conditions *y* ∈ *S ⊆* ℝ ^*l*^, where *S* is a set large enough that the collection of functions {*F* = *f* (., *y*): *y* ∈ *S*} is large enough for determining the one-dimensional distributions of solutions to Eqs. (B.3) in the sense of Eq. (D.1), then solutions to Eqs. (B.3) are unique in distribution.

## Appendix E. Uniqueness of solutions to Eqs. (12)

In this appendix, we establish a duality relation between solutions *X* to the system of Eqs. (12) and the genealogical process *Y*. Applying the result of the previous appendix, we show the uniqueness in distribution of solutions to this system.

Let *X* be a [0, 1]^*k*^-valued solution to the system of Eqs. (12) with *X*(0) = *x* and *Y* the genealogical process with *Y*(0) = *y*. The process *X* has generator

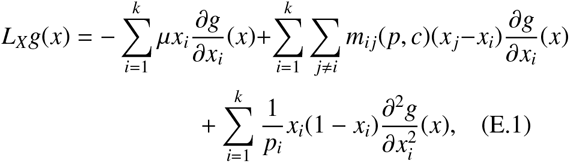

and *Y* has generator

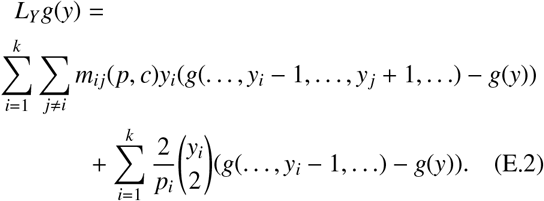

Proceeding informally, for functions *f*: ℝ ^*k*^ × ℝ ^*k*^ → ℝ and *β*: ℝ ^*k*^ → ℝ,

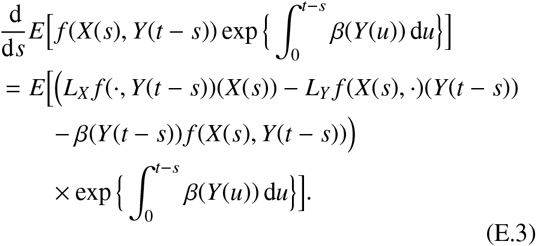

If, then, we can identify judicious choices of *f* and *β* such that

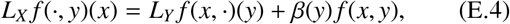

integrating Eq. (E.3) over 0 ≤ *s* ≤ *t* establishes (F). It is easy to verify that taking 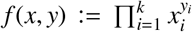 and 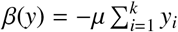 is a suitable choice, and so, assuming that the informal calculation Eq. (E.3) is valid,

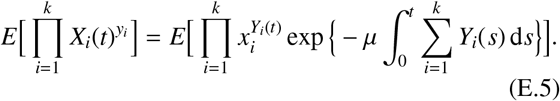

Moreover, since 0 ≤ *X*_*i*_(*t*) ≤ 1 and, because *Y*_*i*_ is a decreasing process, 0 ≤ *Y*_*i*_(*t*) ≤ max_*i*=1,…,*k*_ *y*_*i*_ for all *i* = 1, …, *k, t* ≥ 0, the estimates (G), (H) and (I) hold trivially. In particular, since the genealogical process *Y* can be constructed for any initial condition *y* ∈ ℕ ^*k*^, we have derived an explicit formula for all mixed moments of *X*(*t*), *t* ≥ 0, and, thus, the one-dimensional distributions of all [0, 1]^*k*^-valued solutions *X* to Eqs. (12) with initial condition *X*(0) = *x* coincide. It follows that solutions are unique in distribution.

It remains to justify the informal differentiation of the expectation. We loosely follow the more general arguments of Ethier and Kurtz (1986), pp. 192-195; restricting the analysis to the special case of our setting, the simplicity of the arguments is not lost to abstraction. Setting

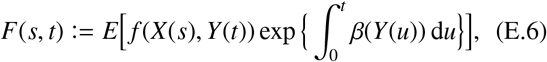

it is clear that

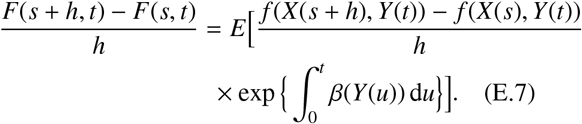

For all test functions *g*, 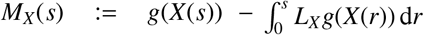 is a martingale (see Theorem 1.6 of Durrett (1996), Chapter 7) and, in particular, we may therefore express the right hand side of Eq. (E.7) as

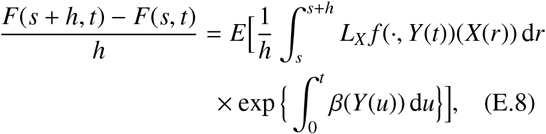

by conditioning on (*Y*(*u*))_*u*≤*t*_ and using that *X* is a process independent of *Y* and so

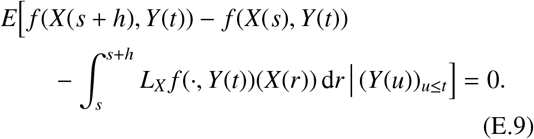

We proceed to pass to the limit *h* → 0. With our choice of *β*, the exponential term inside the expectation on the right hand side is clearly bounded by one; furthermore, by the mean value theorem, a.s.,

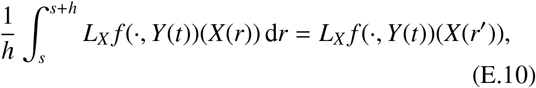

for some *r* ′ with |*r* ′ − *s*| *<* |*h*|, and so, if there exists some constant Γ *<* ∞ such that

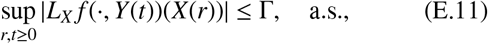

then, by Lebesgue’s dominated convergence theorem, passing to the limit *h* → 0 under the expectation on the right hand side of Eq. (E.8) is valid, and

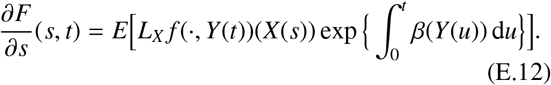

Since, as above, 0 ≤ *X*_*i*_(*t*) ≤ 1 and 0 ≤ *Y*_*i*_(*t*) ≤ max_*i*=1,…,*k*_ *y*_*i*_ for all *i* = 1, …, *k, t* ≥ 0, it is straight-forward to verify the estimate in Eq. (E.11).

Proceeding in the same vein, for *h >* 0 we can write

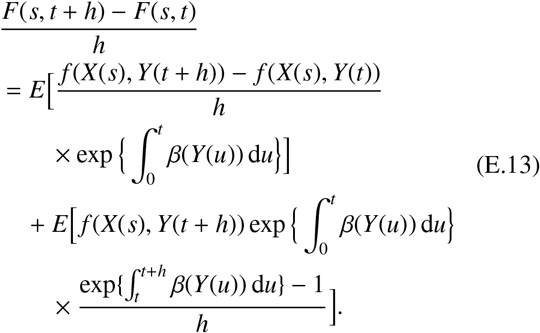

For all test functions *g*, 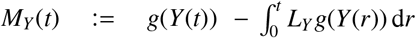 is a martingale. Therefore, by conditioning on (*Y*(*u*))_*u*≤*t*_, the first expectation on the right hand side of Eq. (E.13) may be written as

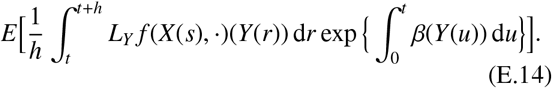

Turning now to the terms in the second expectation on the right hand side of Eq. (E.13), with our choice of the function *β*,

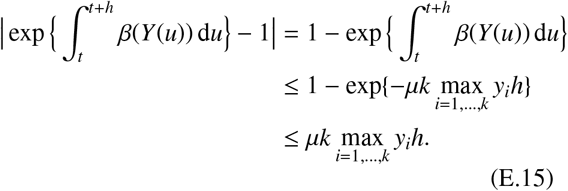

By an analogous argument for the case *h <* 0, and since a.s. the process *Y* does not jump at time *t*, we can apply Lebesgue’s dominated convergence theorem and pass to the limit *h* → 0 in Eq. (E.13) to obtain

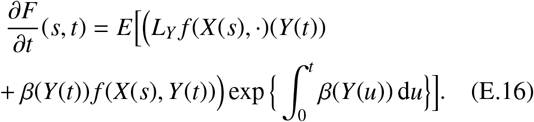

By Eqs. (E.4) and (E.12), it follows that

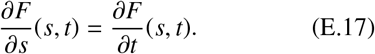

Finally, we note that for *T >* 0,

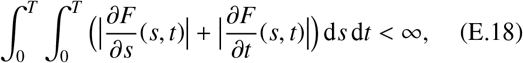

and so, by the Fubini–Tonelli Theorem, exchanging the order of integration yields that

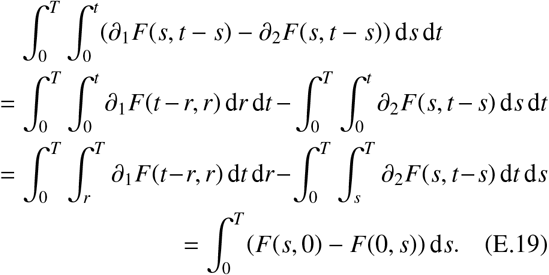

Differentiating with respect to *T*, and using Eq. (E.17), we have verified Eq. (E.5).

## Appendix F. Proof of the diffusion approximation

We will verify the conditions of the Stroock–Varadhan Theorem to show that *X*^*N*^ *⇒ X*, where *X*^*N*^ is defined in Eq. (11) and *X* is the solution of the system of Eqs. (12). Condition (B’), and thus (B), was verified by establishing a duality relation with the genealogical process in the previous appendix.

Let *Q*^*N*^ be defined as in Eq. (C.1); then, since there are *N* individuals each independently entering a reproductive phase at rate *N*, it is clear that *Q*^*N*^ (*x*, ℝ ^k^\{*x*}) ≤*N*^2^ and thus condition (A) is trivially satisfied.

Let 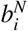 be defined as in Eq. (C.3). By considering each possible outcome of an individual’s reproductive phase,

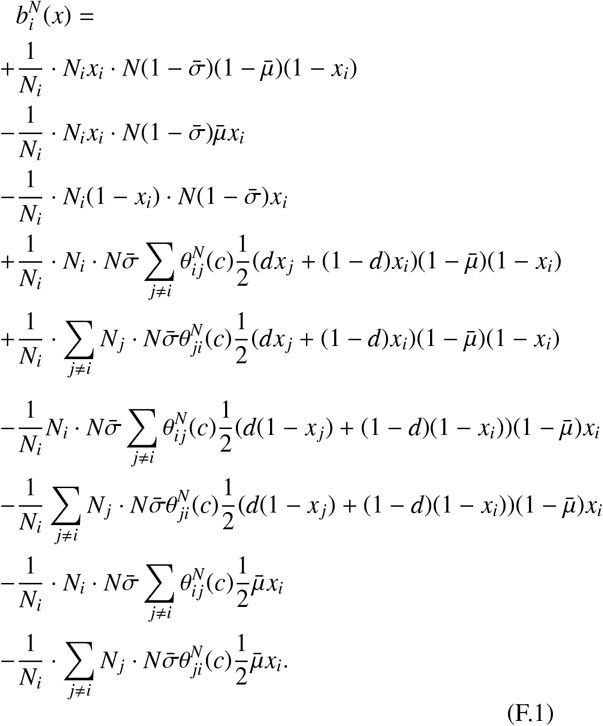

The first term on the right hand side covers the case that the proportion of type *i* individuals carrying the reference type *a*_1_ at the neutral marker *l* increases by 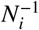 during an asexual reproductive phase; this is the case only if one of the *N*_*i*_ *x*_*i*_ type *i* individuals carrying *a*_1_ enters an asexual reproductive phase (at rate 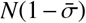 for each individual), there is no mutation (with probability 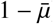 and one of the type *i* individuals not carrying the reference type *a*_1_ at the neutral marker is replaced (with probability 1−*x*_*i*_).

The second and third terms on the right hand side cover the case that the proportion of type *i* individuals carrying the reference type *a*_1_ at *l* decreases by 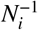 during an asexual reproductive phase; this happens either if one of the *N*_*i*_ *x*_*i*_ type *i* individuals carrying *a*_1_ enters an asexual phase (at rate 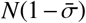), there is a mutation (with probability 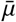), and one of the same group is replaced (with probability *x*_*i*_), or if one of the *N*_*i*_(1− *x*_*i*_) type *I* individuals not carrying *a*_1_ enters an asexual phase (at rate 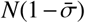 and one of the *a*_1_-carriers is replaced (with probability *x*_*i*_). Note that whether a mutation occurs or not during this latter phase is immaterial since we have assumed there can be no backwards mutation.

The fourth and fifth terms on the right hand side cover the case that the proportion of type *i* individuals carrying the reference type *a*_1_ at *l* increases by 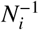 during a sexual reproductive phase in which there is no mutation at *l*. The fourth term corresponds to the first case, in which one of the *N*_*i*_ type *i* individuals enters a sexual reproductive phase (at rate 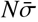) and, for some *j* ≠ *i*, successfully finds a mate of type *j* (with probability 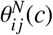 then, with probability (*dx*_*j*_ + (1 − *d*)*x*_*i*_)*/*2, the offspring inherits the mating type *i* and the reference type *a*_1_ at the locus *l*; there is no mutation (with probability 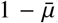) and the replaced individual does not carry *a*_1_ (with probability 1 − *x*_*i*_). The fifth term is derived similarly, except the focal parent is of any type *j* ≠ *i* and the chosen mate is of type *i*. (Here, as in Section 1, we call the individual that enters a sexual reproductive phase the focal parent.)

The sixth and seventh terms cover the case that a type *i* individual carrying *a*_1_ is replaced during a sexual reproductive phase and there is no mutation. These may be derived similarly to the fourth and fifth terms. (Note that in order for a type *i* individual to be replaced, the new individual must be type *i*.)

Lastly, the eighth and ninth terms correspond to the case that, during a sexual reproductive phase, a type *i* individual carrying the reference type *a*_1_ at *l* is lost due to being replaced by a mutant type. The eighth term is the case that one of the *N*_*i*_ type *i* individuals enters a sexual reproductive phase (at rate 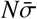) and for any *j* ≠ *i* finds a compatible mate (with probability 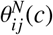), the progeny inherits the mating type from the type *i* parent (with probability 1*/*2), there is a mutation (with probability 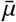), and a type *i* individual which carries *a*_1_ is replaced (with probability *x*_*i*_). The ninth term is derived similarly, except the focal parent is of any type *j* ≠*i* and the chosen mate is of type *i*.

Recall from Eqs. (7)-(8) that, in the regime of rare sex and rare mutation, 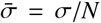 and 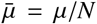. The first three terms sum to

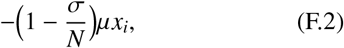

the subsequent four terms sum to

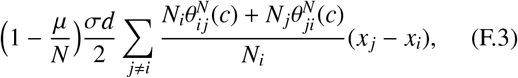

and the final two terms sum to

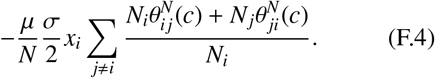

Setting

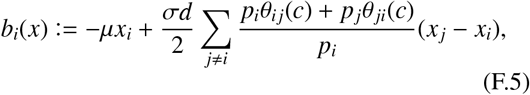

it is straightforward to show that 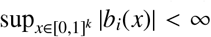 and

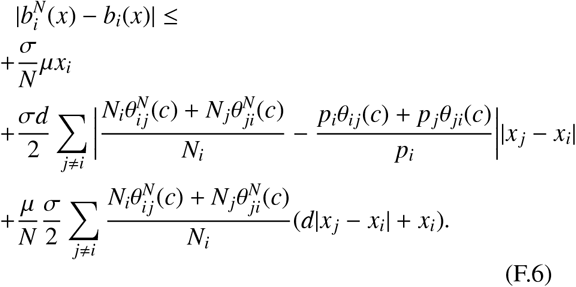

Since we may bound 0 ≤ *x*_*i*_ *≤* 1, it follows that condition (D) is satisfied.

Let 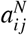 be defined as in Eq. (C.2). By considering each possible outcome of an individual’s reproductive phase, following the calculation of 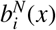 (*x*) exactly,

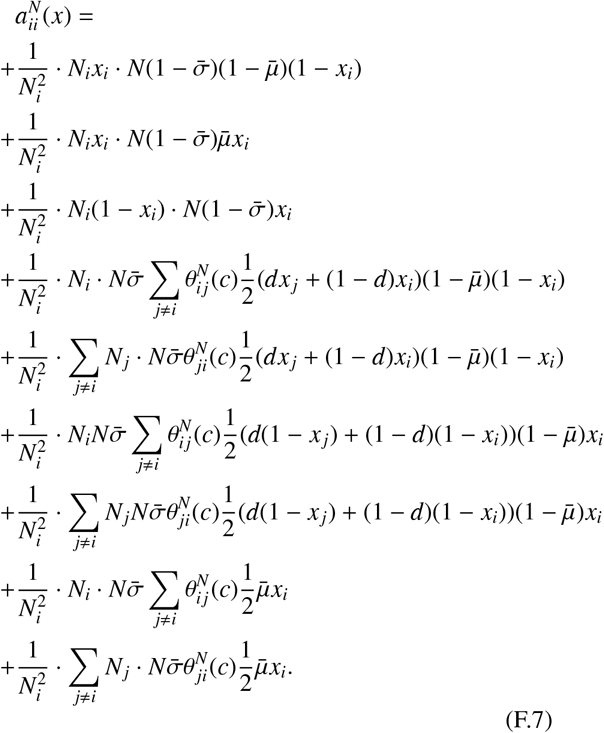

The first three terms sum to

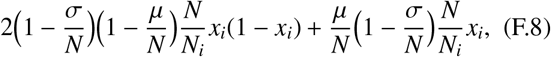

the subsequent four terms sum to

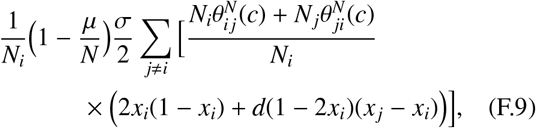

and the final two terms sum to

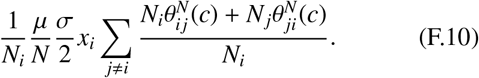

Setting

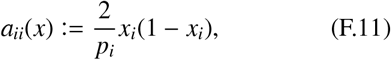

it is straightforward to show, as with the case of the sequence 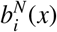 and its putative limit *b*_*i*_(*x*), that condition (C) is satisfied for *i* = *j*. Considering now *i* ≠ *j*, since there can be at most one replacement of an individual during any one reproductive phase, and the new individual is always of the same mating type as the individual that is replaced, we note that there cannot be simultaneous changes in the proportion of type *i* and type *j* individuals carrying *a*_1_ at *l*. In particular, 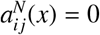 for all *i* ≠ *j* and condition (C) is satisfied with *a*_*i j*_(*x*) = 0.

Let *γ*^*N*^ be defined as in Eq. (C.4). It follows from Jensen’s inequality that

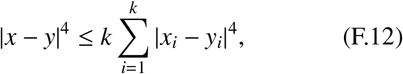

and thus

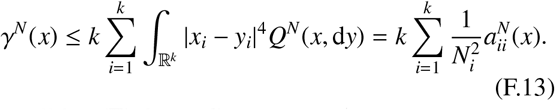

Condition (E) then follows from (C).

## Appendix G. Proof of the approximation of genealogies

Since the dynamics of the continuous-time Markov chain *Y*^*N*^ are described by a bounded Markov generator, it is sufficient to check the convergence of the individual rates.

Consider first a sample of two type *i* individuals in a large population of size *N*. Let us determine how far back in time we have to look until we see an asexual reproductive phase from which our focal individuals were the two outputs (the surviving parent and its progeny). Looking backwards-in-time, either of the ancestral lines we are tracing encounters an asexual phase where it is the parent at total rate 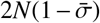 this event concludes by replacing the second ancestral line with probability 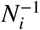. Thus, we encounter a common ancestor (for the allelic type at *l*) after an exponentially distributed amount of time with rate parameter

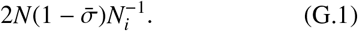

This reproductive phase concludes without mutation with probability 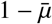, in which case we say that the two individuals coalesce, and with a mutation with probability 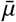. Passing to the limit *N* → ∞, the two individuals coalesce at total rate

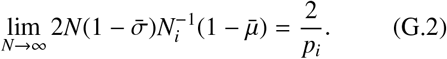

Now let us proceed to the more general setting, considering a sample of *n* individuals from a population of size *N* ≫ *n*. Denote by *n*_*i*_ the number of this sample who are of type *i*. In particular, 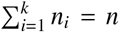. We will establish the genealogy of the sample at the locus *l* in the large population limit *N* → ∞.

We begin by thinking about what happens to the genealogy when we encounter an asexual reproductive phase in the total population. Let us first restrict our attention to the case that the reproductive phase concludes without mutation. These events occur at total rate 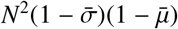, so only events of probability of order 1*/N*^2^ or greater will be seen after passing to the limit. With probability *n*_*i*_*/N*. (*n*_*i*_ − 1)*/N*_*i*_, the asexual phase involves two type *i* ancestral lines in our sample. In this case, the two individuals are said to coalesce. In the limit, this happens then at total rate

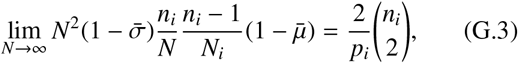

and, after this event, we need only continue to trace *n*_*i*_ − 1 ancestral lines in order to establish the genealogy of our original sample. We must also consider the case that either, or both, of the ancestral lines involved during an asexual phase are not in our sample of *n* individuals. Although this happens at total rate

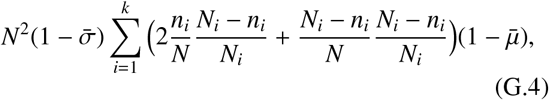

which blows up as *N* → ∞, we still continue to need to trace *n*_*i*_ ancestral lines of type *i* and thus do not see any change in the observed genealogical history.

We now consider the effect on the genealogy when we encounter sexual reproductive phases; again, we begin by restricting our attention to those which do not conclude with a mutation. These events occur at total rate 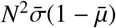. We thus need only consider events of probability of order 1*/N* or greater, since only these will be retained after passing to the limit. Consider when we will encounter a sexual event where one of the sampled *n*_*i*_ type *i* individuals inherits its allelic type from a type *j* individual. This occurs with probability

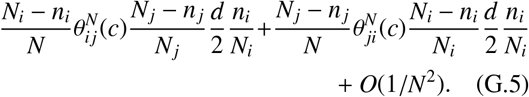

Here, the first term comes from the case where the focal parent is of type *i* but not in our sample of ancestral lines; it successfully finds a type *j* mate; this type *j* mate is also not in our sample; the progeny inherits the mating type from the type *i* individual and the allelic type from the type *j*; and the replaced type *i* individual is in our sample. Another possibility is that the focal parent is of type *j* and the chosen mate is of type *i*, corresponding to the second term in the sum. It would also be the case if either, or both, of the parents are in our sample, but this event is of probability *O*(1*/N*^2^) and thus neglected. Thus, after passing to the limit, we need to continue to trace one fewer type *i* ancestral line and one further type *j* ancestral line at rate

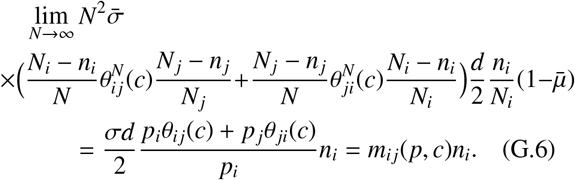

Sexual phases which conclude with the replacement of an ancestral line not in our sample, although possibly occurring with *O*(1) probability, are not seen when tracking just the number of ancestral lines needed to establish the genealogy of our original sample of *n* individuals.

Let us turn now to events which also involve a mutation. In order to observe a change in our genealogical process, the progeny of the event must be in our sample. This occurs with probability *n*_*i*_*/N*_*i*_ if the inherited mating type is *i*. Thus, a reproductive phase is sexual, includes a mutation, and is actually observed in our sample with probability of order 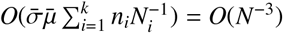. Similarly, a reproductive phase is asexual with a parent in our sample, includes a mutation, and has progeny in our sample with probability of order 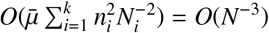. Since events occur at total rate *N*^2^, all such events may be neglected. Turning to asexual phases with parents not in our sample, we observe a mutation when the ancestral line we are tracing is the progeny of an individual not in our sample and a mutation occurs; a type *i* ancestral line experiences a mutation at *l* total rate

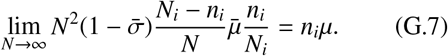

## Appendix H. Identity by descent in a population with *k* self-incompatible mating type classes

In this section we derive a system of equations for the probability that two individuals are identical by descent; our system is specialised to and subsequently solved in the case that *p*_*i*_ = 1*/k* for all *i* = 1, 2, …, *k*.

To ease notation we will write *m*_*i j*_(*p, c*) ≡ *m*_*i j*_. Conditioning on the first coalescence, ‘migration’ or mutation event as we trace ancestral lineages backwards in time,

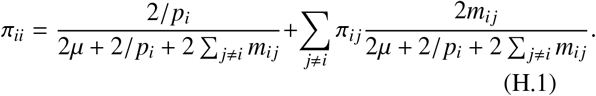

For *i* ≠ *j*,

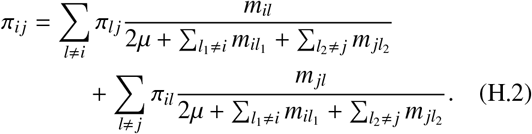

Tractable solutions to this system of equations do not exist in general. Specialising to the case that *p*_*i*_ = 1*/k*, writing *π*_*i j*_ *≡ π*_12_ =: *π*_*d*_ if *i* ≠ *j, π*_*i j*_ *≡ π*_11_ =: *π*_*s*_ if *i* = *j*, and 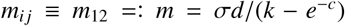, this system of equations may be rewritten as

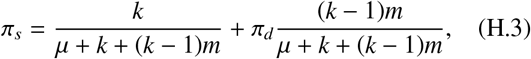

and

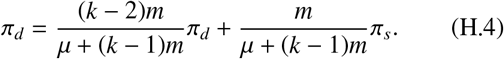

A simple rearrangement of Eq. (H.4) yields that

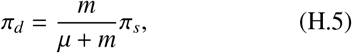

and subsequently substituting this into Eq. (H.3) gives us that

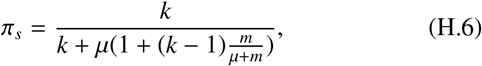

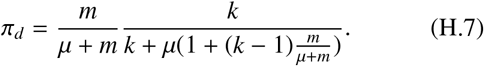

## Appendix I. Identity by descent in mixed self-incompatible and unisexual population

In this section we derive a system of equations for the probability of identity at the neutral marker *l* for pairs of individuals chosen from a population consisting of *k* − 1 SI mating type classes of equal size, and a unisexual class consisting of a proportion *α* of the total population. Let the probabilities 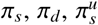 and 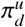 be as defined in Section 2.2.5 above.

We use the limiting genealogical process to compute the probabilities of IBD. Two individuals will be identical by descent if they coalesce before either experiences a mutation.

Two lineages of the same SI type coalesce at rate 2(*k* − 1)*/*(1 − *α*) and two individuals of the unisexual type coalesce at rate 2*/α*. Conditioning on the first event, which may be a coalescence, a mutation or a migration,

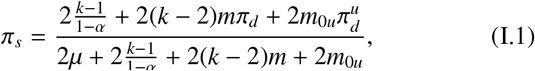

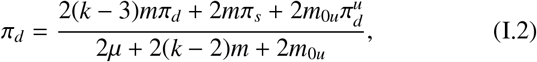

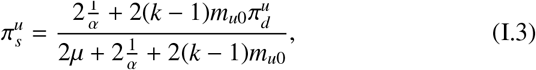

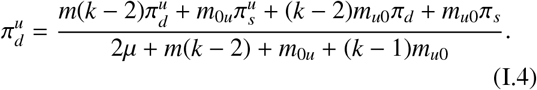

We now consider the case that *α* = 1*/k* and *c* = ∞ in this case, as discussed in the main body of the paper, symmetry between the SI and unisexual classes is introduced and

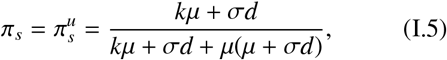

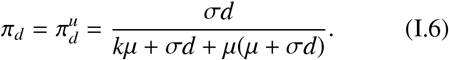

Let us now turn to computing *ϕ* and *ϕ*^*u*^, the probability of mating with an individual distinct at the neutral marker for SI and unisexual individuals respectively. The symmetry does not carry over to these quantities because the unisexual types retain the ability to mate with themselves. The quantities are straightforward to compute:

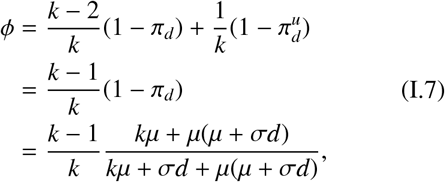

and

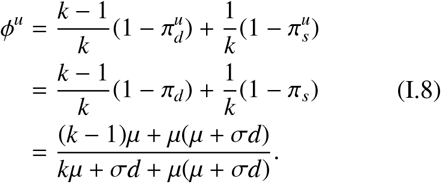

